# Visual working memory representations of naturalistic images in early visual cortex are not sensory-like

**DOI:** 10.64898/2025.12.11.693774

**Authors:** Leonardo Pettini, Carsten Bogler, Karla Matic, Kai Görgen, Christian F. Doeller, John-Dylan Haynes

**Author notes:** Correspondence concerning this article should be addressed to Leonardo Pettini, Address: Charité - Universitätsmedizin Berlin, Bernstein Center for Computational Neuroscience, Charitéplatz 1, 10117 Berlin. No conflicts of interest to declare.

## Abstract

For a long time, working memory was considered to reflect sustained activity in prefrontal and parietal areas. This has been challenged by the “sensory recruitment model” according to which sensory areas are involved in maintaining working memory representations. The main support for this comes from the finding that working memory contents can be decoded from early sensory areas during memory delay periods. However, the availability of stimulus-related information in sensory regions does not reveal whether this information is encoded in a sensory format, i.e. in the same way as sensory stimuli. Alternatively, working memory contents could be encoded in independent formats or even potentially undergo dynamic changes throughout the delay. Here, we test this question directly by requiring participants to briefly memorise naturalistic object-scenes. We directly compared the way early visual regions encoded the contents during perception and working memory maintenance. As in previous studies, we found robust stimulus-related information in early visual cortex throughout the delay. However, despite continuously robust information, the encoding of memory contents in the later delay period did not occur in the same format as during pure perception. Thus, despite the fact that sensory areas have working-memory-related information, the format of this information is not strictly sensory-like throughout the delay.

## 1 Introduction

Working memory supports the short-term maintenance of visual information in the absence of sensory input (Fuster & Alexander, 1971; Miller & Cohen, 2001; Eriksson et al., 2015; Baddeley, 2020), and enables the temporal processing and integration of perception to guide behaviour (van Ede & Nobre, 2023). The localisation of working memory function in the brain has been debated. The prefrontal cortex (Jacobsen, 1936; Fuster & Alexander, 1971; Kubota & Niki, 1971; Funahashi et al., 1989; Miller et al., 1996), along with temporal and parietal association areas (for a review, see Constantinidis and Procyk (2004)) has long been regarded as central to working memory. Early non-invasive human neuroimaging studies corroborated this view, demonstrating similar prefrontal and parietal activations (Jonides et al., 1993; D’Esposito et al., 1995; Doyon et al., 1996; Courtney et al., 1997; Courtney et al., 1998; Wager & Smith, 2003).

This account began to be challenged by the idea that working memory might rely on encoding in sensory regions (Curtis & D’Esposito, 2003; Pasternak & Greenlee, 2005). An influential line of evidence came from functional magnetic resonance imaging (fMRI) studies using Multivariate Pattern Analysis (MVPA), which demonstrated that stimulus-specific information could be decoded from early visual cortex during working memory maintenance (Harrison & Tong, 2009; Serences et al., 2009; Christophel et al., 2012). These results supported the proposal that sensory circuits might actively maintain mnemonic information across working memory delays, besides their involvement in bottom-up sensory processing (Curtis & D’Esposito, 2003; Pasternak & Greenlee, 2005; Postle, 2006; D’Esposito, 2007; D’Esposito & Postle, 2015). This framework became known as the sensory recruitment model (Awh & Jonides, 2001; Serences et al., 2009).

A critical issue is whether sensory and memory representations share the same representational format, yet this question is rarely tested directly. Note that, despite being in early visual areas, the encoding of working memory contents could also occur in independent formats, which would for example have the advantage that it would be easier to maintain a separation between bottom-up sensory processing and working memory representation (Rademaker et al., 2019; Libby & Buschman, 2021; Degutis et al., 2025). Evidence supporting sensory-like working memory codes comes from studies in which classifiers or encoding models trained on perceptual data (e.g. from the encoding period of the working memory task or an independent localizer) successfully generalise to predict working memory content in early visual cortex (Harrison & Tong, 2009; Albers et al., 2013; Rademaker et al., 2019; Iamshchinina et al., 2021; Vo et al., 2022). However, there is also some evidence that working memory codes might not be sensory. For example, memory decoding is weaker when it is based upon sensory-trained classifiers compared to memory-trained classifiers in the afore-mentioned studies, implying that the representational format changes between perception and maintenance. In other cases, cross-decoding effects are not even statistically significant (Lorenc, 2020). Recently, Duan and Curtis (2024) used special stimuli to compare the cross-decodability of sensory and memory codes. While sensory representations in early visual cortex were specific to each modulated grating, working memory representations generalised across them, consistent with working memory representations being an abstraction of sensory features (Kwak & Curtis, 2022). Finally, there is evidence that working memory codes might be maintained dynamically throughout the delay phase, as proposed by electrophysiology (Meyers et al., 2008; Stokes et al., 2013; Spaak et al., 2017; Meyers, 2018) and fMRI work (Sreenivasan et al., 2014; Li & Curtis, 2023; Degutis et al., 2025), undergoing a reformatting from sensory-like to memory-specific representations. Another less investigated question is whether the cross-decodability between sensory and memory representations holds only for simple geometric stimuli or also for more complex, naturalistic stimuli. Most evidence for sensory recruitment comes from studies using low-dimensional visual stimuli such as orientations (Harrison & Tong, 2009; Pratte & Tong, 2014; Ester et al., 2015), colours (Serences et al., 2009), motion parameters (Riggall & Postle, 2012; Emrich et al., 2013), abstract shapes (Christophel et al., 2012) and flow-field patterns (Christophel & Haynes, 2014). Despite a growing interest in more naturalistic working memory processes (Brady et al., 2019; Xu, 2023; Bates et al., 2024; Adam et al., 2025), sensory recruitment for naturalistic images has only rarely been addressed (e.g. Xu, 2023).

Here, we address this gap and directly test whether delay-period activity in early visual cortex reflects sensory-like representations. We conducted an fMRI study involving two sessions. In the first session, participants performed a delayed-match-to-sample task with a custom stimulus set consisting of synthetically generated, complex, naturalistic object-scenes (Pettini et al., 2025). Our targets and foils were perceptually very similar and closely psychophysically characterized. This custom-designed set of naturalistic images allows to closely titrate the similarity between targets and foils and thus ensure that successful performance required that subjects memorise the fine-grained details of the object-scenes rather than coarse or categorical differences. Moreover, since target and foils differed in overall similarity, participants had to attend to each image as a whole, rather than focusing on specific sections. The working memory task was therefore more challenging than it would have been if they had to identify fundamentally different categorical stimuli. Also, because targets and foils are only slightly different realizations of the same scenes, verbal rehearsal is of limited use. In the second session, participants performed a purely perceptual task with no memory component, which provided us with an independent sensory training set for multivariate classification.

We found robust stimulus-specific classification in early visual cortex regarding the natural scenes. First, training and testing on the purely perceptual task revealed robust information in early visual cortex during bottom-up perception. Second, training and testing on the memory task also revealed robust information throughout the entire delay. This decodable information when training and testing on the memory experiment was highest in the early visual cortex, even for the naturalistic stimuli used in our paradigm. Third, however, the information that could be extracted by training on the perceptual task and testing on the memory task did not last throughout the delay. Thus the sensory-to-memory generalisation was robust but only for the initial period of the delay period. In the later phase of the delay, there was substantial information for classifiers trained on memory, but not for classifiers trained on perception. This pattern was consistent across multiple classification algorithms, correlational measures and regions of interest.

## 2 Methods

Data were collected within a larger study investigating the effects of repeated experience with a stimulus on its working memory maintenance (preregistered under https://osf.io/tyxh8).

### 2.1 Participants

We recruited 53 participants through flyers, university participant databases, and personal referrals. Eligibility criteria included being healthy, aged 18–40, right-handed, with normal or corrected-to-normal vision, and no history of neurological or psychiatric disorders. All participants were screened for MRI compatibility using standard procedures at the Berlin Center for Advanced Neuroimaging. Participants provided informed consent prior to participation and received a total monetary compensation of 72 Euro. The study was approved by the ethics committee of the Humboldt-Universität zu Berlin and conducted in accordance with the Declaration of Helsinki. Of the initial 53 participants, we excluded 11 participants across the two sessions due to: excessive head motion (1), hardware or software issues (3), failure to understand the task or to follow the instructions (2), incidental findings (1), withdrawal of consent (3), and failure to perform the task due to marked drowsiness (1). Our final sample included 42 participants (23 female, 19 male; M = 27.1 years, SD = 5.1).

### 2.2 Stimuli

We used a delayed match-to-sample task in combination with naturalistic images that were custom-designed to prevent reliance on verbal strategies alone and required participants to encode fine-grained perceptual differences for discriminating between target images and foils. To achieve this, target and foil images shared the same semantic identity but differed in their specific visual features. Importantly, we manipulated the overall target-foil similarity rather than single perceptual features (e.g. colour alone), so that the task would require participants to attend to multiple visual details.

Specifically, our custom stimulus set consisted of 63 synthetically generated object-scenes (Figure 1a), subsampled from a larger set (Pettini et al., 2025). Every object-scene represents a single object positioned centrally within a coherent context (e.g., a cat on a carpet), spanning three natural (animals, plants, landscape elements) and three artificial categories (items, vehicles, buildings). For each object-scene, four perceptual variations of increasing dissimilarity but invariant semantic identity were selected using the procedure described in Pettini et al. (2025). The first variation served as the target image, while the remaining three functioned as foils with decreasing perceptual similarity to the target (Figure 1b), yielding three difficulty levels (Easy, Medium, Hard). An additional 10 object-scenes were selected for the behavioural training session outside of the scanner.

**Figure 1:**
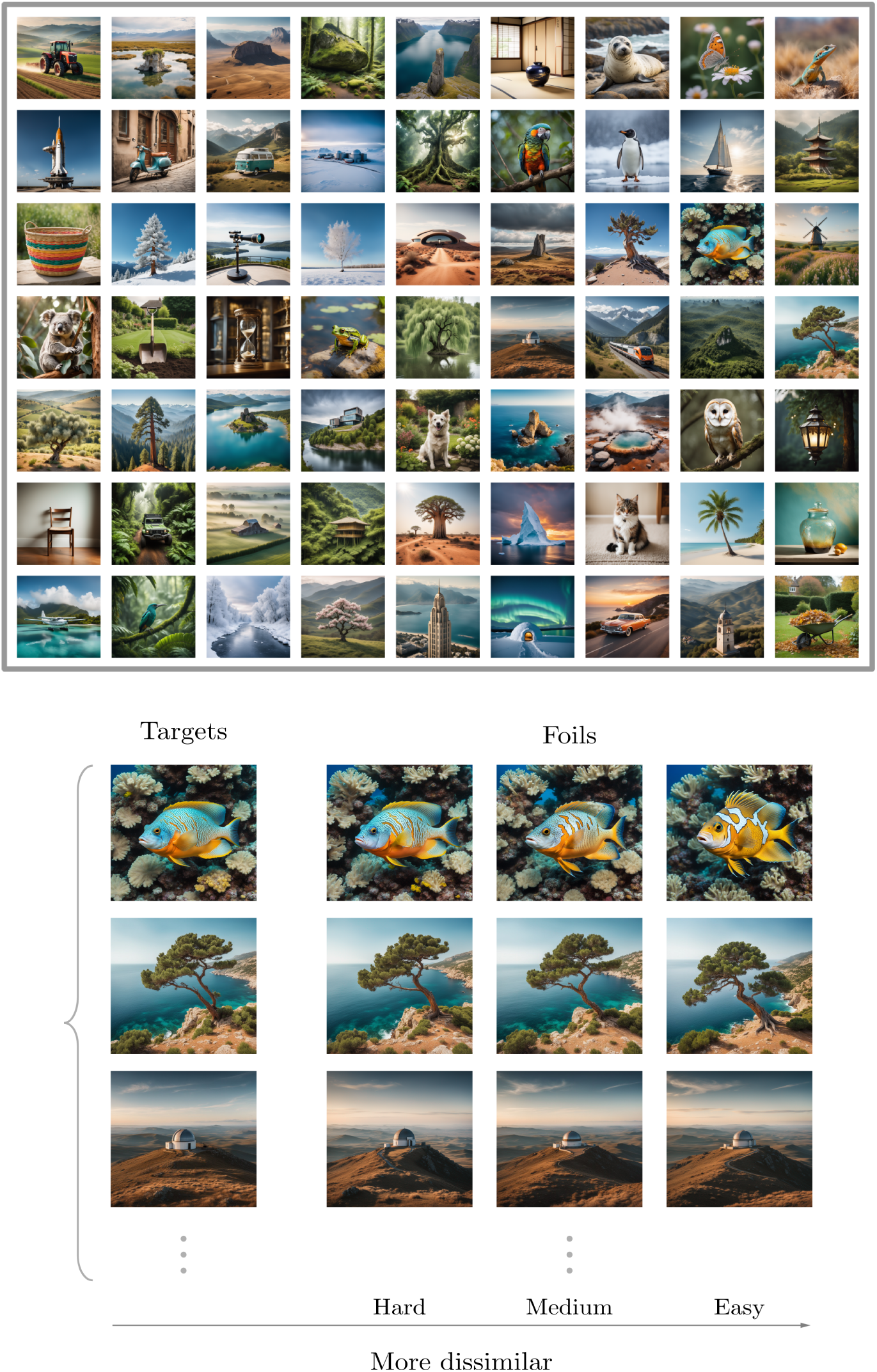
Overview of the experimental stimuli and task. (a) As targets, we used 63 object-scenes from a novel, synthetically generated set of naturalistic images (Pettini et al., 2025). (b) For each object-scene, we selected four variations (one target and three foils of increasing dissimilarity). The scaled perceptual dissimilarity was validated in an online crowdsourcing task (*N* = 1113), where participants judged the perceptual similarity between the object-scene variations (Pettini et al., 2025).

All images were shown on a uniform grayscale background. A central bull’s-eye with crosshairs (Thaler et al., 2013) served as the fixation target, which remained visible throughout the run and was overlaid on every image during presentation. The experiment was programmed and run using the Python package PsychoPy v2024.2.1 (Peirce et al., 2019). The stimuli were displayed on an MRI-compatible screen (52 cm × 39 cm, 1024 × 768 px resolution) positioned at the rear of the scanner bore, which participants viewed through a mirror mounted on the head coil. The viewing distance from the eyes to the centre of the screen was 158 cm and the stimuli subtended a visual angle of approximately 7 degrees.

### 2.3 Task Design

The experiment included two scanning sessions (Figure 2): in Session 1, participants performed a delayed match-to-sample task with repeated and non-repeated targets (”memory task”); Session 2, on the other hand, was a perceptual viewing task that provided an independent training set for classification analyses (”perceptual task”).

**Figure 2:**
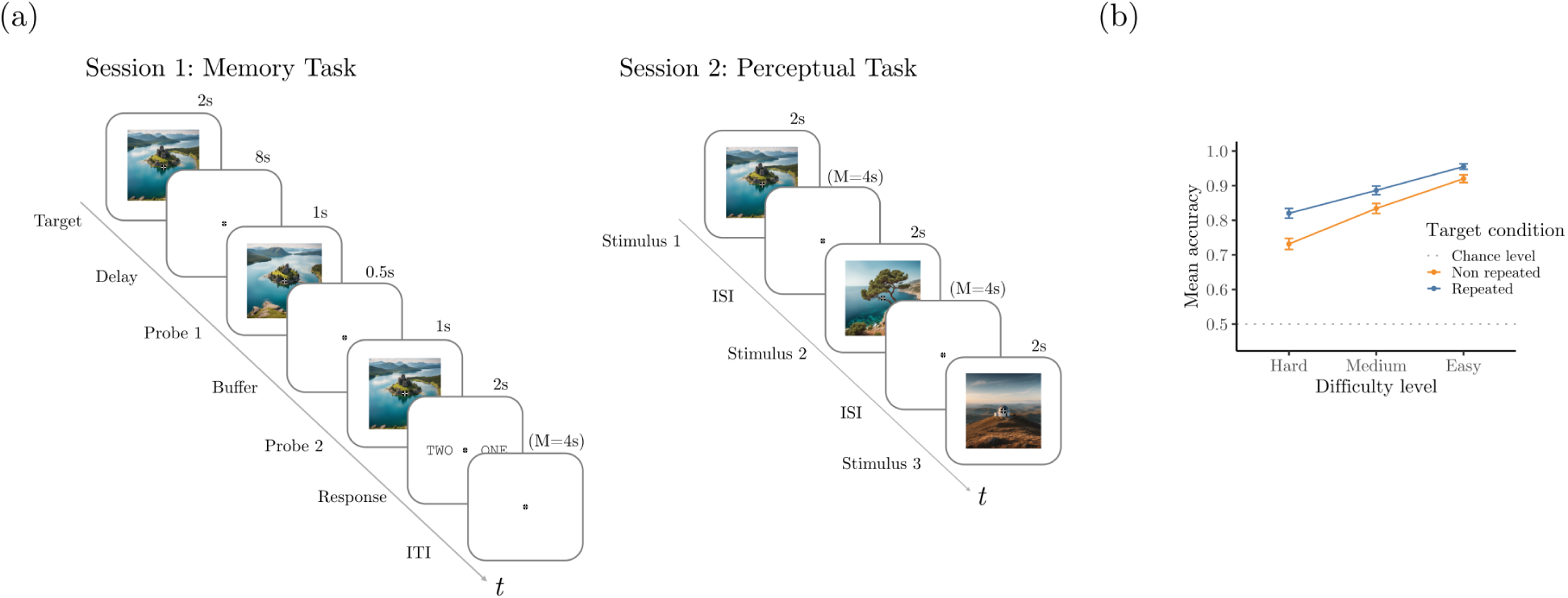
(a) Schematic representations of the two tasks. Session 1 (memory task): each trial included stimulus presentation (2 s), an 8 s delay, a first probe (1 s), a 0.5 s buffer, a second probe (1 s), a response window (2 s), and a jittered intertrial interval (mean 4 s). Participants performed the task with 9 repeated and 54 non-repeated targets. Repeated targets were shown once in each of the 6 blocks. Session 2 (perceptual task): participants viewed all images from Session 1 for 2 s each, separated by jittered interstimulus intervals (ISI; mean 4 s). Each object-scene was viewed once per run, 8 times in total. (b) Overall mean recognition accuracy for each difficulty level. Error bars represent ±1 standard error of the mean across participants. Dotted line indicates chance level (50%). More similar target-foil pairs were more difficult to discriminate, leading to lower mean accuracies.

#### 2.3.1 Working Memory Task

The memory task used in Session 1 consisted of six runs of 18 trials each, for a total of 108 trials. Each trial began with the presentation of a target stimulus for 2 seconds, which participants were instructed to memorise throughout a subsequent 8-second delay period. In the delay phase, only the fixation cross was visible on screen. At the end of the delay phase, two probes (one being the target and one the foil) were presented sequentially for 1 second each, separated by a 500-ms interval. Following the offset of the second probe, a response screen displaying the options “one” and “two” was shown for 2 seconds. Participants were instructed to select option “one” if they believed the first probe was the target image and option “two” if they believed it was the second. Responses were given using the index and middle fingers of the right hand via a two-button response box. The spatial position of these two response options (i.e. which of the two was placed on the left or on the right side of the fixation cross) was pseudo-randomised and counterbalanced. The order in which target and foil were presented was also pseudo-randomised and counterbalanced across the experiment. The response screen duration was fixed at 2 seconds, regardless of when the button was pressed. No feedback was provided upon button press. Jittered inter-trial intervals (ITIs) were sampled from a truncated exponential distribution with a mean of 4 s and range of 2-8 seconds. The trial duration from target presentation to the end of the response window was 14.5 seconds, the average trial duration including the ITI was 18.5 seconds. Stimulus assignment to conditions (repeated vs. non-repeated) was pseudorandomised across participants, with the constraint that each object-scene served as a repeated target for six participants across the sample.

Trials were assigned one of three difficulty levels (Easy, Medium, or Hard) based on the similarity between the target and the foil image. Within each run, trial difficulty was balanced across target conditions, with three trials per difficulty level for both repeated and non-repeated targets. Since each repeated target was presented once per run across six runs and there were three difficulty levels, we assigned two trials per difficulty level to each repeated target. We pseudorandomised this assignment with the constraint that all difficulty levels had to be used once for each repeated target image, before they could be used again. This was designed to have a uniform difficulty for each object-scene throughout the experiment.

#### 2.3.2 Perceptual Task

Session 2 was a separate scanning session that was designed to provide an independent estimate of the perceptual representation code of the 63 targets from Session 1. It was conducted not later than 48 hours after the first session. Participants viewed each of the 63 target images once per run across 8 runs, for a total of 504 presentations (8 per stimulus). The stimulus order was randomised within runs. Each stimulus was presented for 2 seconds, and jittered ITIs were drawn from a truncated exponential distribution (mean = 4 s, range = 2–8 s). To keep participants engaged, we included an independent change-detection task: on occasional trials, the fixation cross randomly turned green or red for 1 second after image presentation, and participants responded to the change by pressing the left or right button, respectively.

### 2.4 Experimental Procedure

The two scanning sessions lasted a maximum of two hours each, including non-scanning time (e.g. instructions and breaks between runs). In both sessions, participants first reviewed and signed the consent form, followed by a detailed briefing on the experimental tasks and safety procedures, in accordance with the guidelines of the Berlin Center for Advanced Neuroimaging Charité–Universitätsmedizin Berlin. Before Session 1, they completed a behavioural training session to familiarise themselves with the task before entering the scanner. A separate set of object-scenes was used for this training. Each run of Session 1 lasted 5 minutes and 46 seconds. This total included a 6-second buffer period added at both the beginning and end of each run. No training was conducted at the beginning of Session 2. Each run of Session 2 lasted 6 minutes and 31 seconds, with an additional 6-second interval at both the beginning and end. Participants were instructed to maintain stable fixation on the central fixation target throughout the experiments.

### 2.5 Data Acquisition

MRI data were collected using a 3T Siemens Prisma scanner (Siemens, Erlangen, Germany) with a 64-channel head coil. Two high-resolution T1-weighted MPRAGE structural images were acquired at the beginning of each session (208 sagittal slices, TR = 2400 ms, TE = 2.22 ms, TI = 1000 ms, flip angle = 8°, voxel size = 0.8 mm isotropic, FOV = 166.4 × 240 × 256 mm³). T2-weighted functional images were acquired using a multiband-accelerated EPI sequence (multiband factor = 8; TR = 800 ms, TE = 37 ms, flip angle = 52°, voxel size = 2 mm isotropic, 72 contiguous slices, FOV = 208 × 208 × 144 mm³). For Session 1, we recorded a series of 433 volumes per run (6 runs, 5 min 46 s each). For Session 2, we recorded a series of 489 volumes (8 runs, 6 min 52 s each). At the beginning of each run, we acquired two spin-echo field maps with anterior–posterior (AP) and posterior–anterior (PA) phase-encoding directions to estimate magnetic field inhomogeneities for distortion correction during preprocessing.

### 2.6 Analysis of fMRI Data

#### 2.6.1 Software

Statistical fMRI analyses were conducted using Python (v3.11.5). For neuroimaging data preprocessing, including image loading, masking, signal cleaning, and both first and second level modelling, we used Nilearn (v0.11.1) (Nilearn contributors et al., 2025). Classification analyses were implemented using algorithms from scikit-learn (v1.5.1) (Pedregosa et al., 2011). They were performed both on GLM parameter estimates and raw fMRI time series using linear support vector machines (LinearSVC class) from scikit-learn.

#### 2.6.2 Preprocessing

Raw data were first converted from DICOM to NIfTI format using dcm2bids (v3.1.1), in accordance with BIDS specifications. Preprocessing was carried out with fMRIPrep (v24.1.0rc0) (Esteban, Markiewicz, Blair, Moodie, Isik, Erramuzpe, et al., 2019). T1-weighted anatomical images were corrected for intensity non-uniformity, skull-stripped, segmented into tissue classes, and normalised to the Montreal Neurological Institute (MNI) standard space. A T1-weighted reference image in subject space was computed from the four available scans (two per session). For each BOLD run, fMRIPrep generated a per-run BOLD reference scan, which was then used to estimate and correct for head motion with FSL’s mcflirt (six motion parameters), prior to any spatial or temporal filtering. The reference was co-registered to the subject’s T1-weighted image using FreeSurfer’s mri coreg, followed by FSL’s flirt with boundary-based registration, constrained to six degrees of freedom. Confound time series were extracted from the preprocessed functional data, including framewise displacement (FD) and DVARS (Power et al., 2012), and global signals from CSF, white matter, and whole-brain masks and component-based noise regressors (CompCor) (Behzadi et al., 2007a). For the GLM analyses (see below) we applied spatial smoothing with a 5 mm full-width at half-maximum Gaussian kernel. The full boilerplate output of fMRIPrep detailing all preprocessing steps, software versions and references is available in the Supplementary Methods (Appendix A.2).

#### 2.6.3 Definition of Regions of Interest (ROIs)

We defined the early visual cortex region of interest (EVC) by combining the probabilistic anatomical maps for V1, V2, and V3 from Wang et al. (2015). For each of the three areas, we first created a single continuous bilateral mask in MNI space by combining the left and right hemisphere maps, along with their dorsal and ventral subdivisions. We then used the inverse normalization parameters estimated by fMRIPrep to warp the maps from standard space into each participant’s native space using the Advanced Normalization Tools (ANTs) (Tustison et al., 2021) at 0.8 mm isotropic resolution, and for functional analyses, into the T1-aligned BOLD reference space at 2 mm isotropic resolution. In native space, we thresholded the probability maps at 10 percent, retaining only voxels with at least 10 percent probability of belonging to the target ROI. The average size of the early visual cortex ROI was 6,882 (SD = 873) voxels, with 3,845 (SD = 417) voxels excluded on average due to thresholding. In addition, we conducted additional supplementary and exploratory analyses on other ROIs provided by Wang et al. (2015) to examine the potential contribution of higher visual and parietal areas beyond early visual cortex, such as lateral occipital regions (e.g. LOC) and intraparietal sulcus regions (e.g. IPS). All other maps were processed using the same procedure as early visual cortex. All ROI sizes are shown in Figure S1 (Appendix A.5).

#### 2.6.4 GLM Modelling

For each participant we first estimated two separate GLMs, one for each session. The GLM for Session 1 used separate regressors for the stimulus presentation phase (2 s) and the delay phase (8 s), for each of the 18 trials per each of the six runs (yielding a total of 2 x 108 regressors). We also included six motion parameters (3 translations and 3 rotations) as well as left and right button presses as nuisance regressors. All regressors (except for the motion regressors) were convolved with a canonical SPM Haemodynamic Response Function (HRF). Low-frequency drifts were modelled using a discrete cosine basis set and a high-pass filter cutoff of 0.008 Hz was used. We accounted for temporal autocorrelation in the noise using an autoregressive model of order 1. For Session 2, we again used a trial-wise GLM with separate 2-second event-regressors for each trial coinciding with the stimulus presentation in that trial. We also used six motion parameters (3 translations and 3 rotations) as well as left and right button presses as nuisance regressors. All regressors except the motion regressors were convolved with a standard HRF. All regressors were estimated separately for each of the eight runs. We used the same parameters for removal of low- and high-frequency noise as well for Session 1.

#### 2.6.5 GLM-based Decoding

The single-trial parameter estimates for each trial, run, voxel, and region of interest were then entered into a series of classification analyses.

##### Session2-Session2

For the perceptual task in Session 2 we conducted a GLM-based classification analysis based on single-trial parameter estimates for each of the 63 stimuli (e.g. “cat” or “olive tree”). We assessed classification performance using leave-one-run-out cross-validation, training a linear multiclass SVM (with regularization parameter C=1) on single-trial parameter estimates from seven runs and testing on the held-out run. The classifier employed 63 labels, each representing a different object scene. Thus, chance level for this multiclass classification was 1*/*63 images (≈1.6%). The perceptual task in Session 2 provides an independent and unbiased training set, allowing us to build robust classifiers based on sensory representations.

##### Session2-Session1

We then tested whether these classifiers would generalise from the perceptual task in Session 2 to the working memory task in Session 1. We now trained the linear multiclass SVM (C=1) on parameter estimates from all runs of the perception-only condition in Session 2. We then applied this classifier separately to the parameter estimates for the stimulus presentation and delay periods of Session 1. Classifiers were tested across all six runs of Session 1 with a chance level of 1*/*63 images (≈1.6%). Note that here it was possible to use all available trials (see below) and thus maximize sensitivity.

##### Session1-Session1

We then conducted three GLM-based classification analyses within the memory task in Session 1, training and testing classifiers using parameter estimates from stimulus presentation and delay phase. We trained and tested classifiers using parameter estimates from the same condition (stimulus-stimulus, delay-delay). We also decoded across conditions (stimulus-delay). Importantly, limiting the analysis to the memory session offered a more sensitive test by avoiding cross-session variability. We used leave-one-run-out cross-validation, training linear multiclass SVM (C=1) on GLM parameter estimates from five runs and testing on the held-out run. Note that these analyses were restricted to repeated stimuli, because non-repeated stimuli were presented only once across the experiment and thus could not be used for training and testing across runs. Thus, the chance level for this multiclass classification was 1/9 (≈11%).

##### Statistical analysis of GLM-based classification accuracies

In all analyses, classifier accuracy was first computed for each participant and each cross-validation fold separately. It was then averaged across the 6 (within-Session 1 analysis) or 8 (within-Session 2 and cross-session analyses) cross-validation folds, obtaining a cross-validated performance estimate for each participant. To assess statistical significance, subjectwise accuracies were entered into a second-level one-sample permutation test using a sign-flipping procedure (Nichols & Holmes, 2002; Winkler et al., 2014). For each ROI, we subtracted chance level from each subject’s mean accuracy. If the null hypothesis is correct, the true population mean difference from chance level is zero, and the signs of the differences between individual subject’s mean accuracies and chance are interchangeable. After centering the accuracy values by subtracting chance, we flipped their signs randomly across 10,000 permutations. For each permutation, we computed the obtained group mean difference and compared it to the mean difference observed experimentally. We calculated p-value as (number of permutations with mean accuracy ≥ observed mean accuracy + 1)/(10,000 + 1). When multiple conditions and ROIs were analysed, we controlled for multiple comparisons using the false discovery rate (FDR) correction with the Benjamini-Hochberg procedure (Benjamini & Hochberg, 1995). We used these corrected p-values to assess statistical significance, which was determined as *p <* 0.05.

#### 2.6.6 Time-resolved classification

In order to characterise how stimulus-specific representations evolved over time during the memory task in Session 1, we additionally performed a series of time-resolved decoding analysis based on the raw voxel time series. BOLD time series signals were extracted from all predefined ROIs (see above), linearly detrended and z-standardised on a run-wise basis.

##### Session2-Session1

In a first analysis, we trained SVMs on all available data from Session 2 (see below) and decoded object identity at each timepoint of each trial in Session 1. This analysis aimed to assess the temporal profile of stimulus-specific information across sessions and was conducted separately for each participant. In Session 2, each run provided one sample per label, resulting in a total of 63 labels and a chance level of ≈1.6%. In order to capture the peak of the haemodynamic response to the perceptual stimuli, trial-wise z-scored BOLD signals were averaged within a fixed time window from 4 to 8 seconds after stimulus onset. For each participant, classifiers were trained on 504 samples (63 images × 8 runs) and a feature set corresponding to the number of voxels in the participant’s mask. We then tested each classifier at every timepoint within each trial of Session 1, spanning the stimulus presentation, delay, probe, and response phases. As during training, the BOLD signal was z-standardised independently for each run. For each trial, we extracted a window of 25 consecutive TRs (20 s), time-locked to stimulus onset during the presentation phase. All TRs within the trial time window were assigned the same label, that is the object identity of the target image used in that trial (e.g. “cat”). Classifier predictions and accuracies were calculated at each timepoint. For each participant, we computed average classification accuracy at each timepoint across the full trial window. Finally, we estimated time-resolved accuracy curves with 95% confidence intervals across participants. The same pipeline was repeated using different ROIs.

##### Session1-Session1

We then conducted a time-resolved analysis within Session 1. This analysis was restricted to repeated target images, as non-repeated targets were presented only once per participant and were therefore unsuitable for training and testing a classifier. BOLD signals were extracted from continuous timepoints (TRs), linearly detrended, and z-standardised independently for each run. For each trial, we defined a fixed window of 25 TRs, time-locked to stimulus onset (TR = 0). As the TR was 0.8 seconds, this window spanned 20 seconds and covered the full trial duration. The window included the stimulus presentation (0-2 s), delay period (2-12 s), probe presentation (12-14.5 s), and response phase (14.5-16.5 s), as well as 3.5 seconds following trial offset (see Methods for task timing). For each ROI and participant, we trained separate linear SVMs at each individual timepoint within this 25-TR window. Training and testing occurred at the same temporal position across each trial. We used leave-one-run-out cross-validation: in each fold, data from one run were held out for testing while the remaining five runs were used for training. As each repeated image was presented once per run, the classifier was trained with five samples per label and tested on the sixth. Nine labels were included per participant, resulting in a theoretical chance level of 1/9 (≈11%). Classification accuracy for each timepoint was computed per fold and subsequently averaged across folds within participants. The final result was a decoding accuracy time course for each participant.

##### Statistical Analysis of Time-Resolved Classification

To determine at what time-points the accuracy time courses were significantly above chance, we used a non-parametric, cluster-based permutation procedure, which controls for multiple comparisons (Bullmore et al., 1999; Maris & Oostenveld, 2007a; Groppe et al., 2011). For each ROI, we computed a one-sample, one-sided t-statistic at each timepoint to test if classification accuracy was significantly above chance. We then grouped contiguous timepoints that exceeded a predefined cluster-defining threshold (t-value corresponding to *p <* 0.05). The sum of the t-values within each cluster (the cluster “mass”) served as our test statistic. To generate a null distribution for this statistic, we performed 10,000 sign-flip permutations across participants. Finally, cluster-wise p-values were calculated by comparing the mass of each observed cluster against this null distribution, and these p-values were assigned to all timepoints within their respective significant clusters.

#### 2.6.7 Temporal Generalization Analysis

To assess the temporal generalisation of brain activity patterns over the course of a trial and across different ROIs, we conducted a temporal generalisation classification analysis in Session 1 (Anders et al., 2011; King & Dehaene, 2014). Following the same preprocessing steps as in the time-resolved analysis within Session 1, we extracted BOLD signals from continuous TRs, linearly detrended them and z-standardised them independently for each run. For each trial, we again analysed a fixed time window spanning 25 TRs (20 s), time-locked to the onset of stimulus presentation. However, rather than training and testing at the same timepoint, we now trained separate linear SVMs at each individual timepoint and tested each classifier across all timepoints within the trial. This cross-temporal decoding approach was applied separately for each ROI and participant, yielding a temporal generalisation matrix for each combination.

##### Statistical Analysis of the Temporal Generalization Matrices

To assess statistical significance and control for multiple comparisons in the temporal generalisation analysis, we used cluster-based permutation testing (Maris & Oostenveld, 2007b). First, we subtracted the theoretical chance level from the classification accuracies at each training–testing timepoint pair for each participant. We then computed a one-sample t-statistic at each pair of timepoints across participants, resulting in a group-level t-map. A threshold was applied to the t-map corresponding to a one-tailed test at *α* = 0.05 (df = 42). Clusters were defined as contiguous sets of timepoints where the t-values exceeded this threshold, so that only elements with evidence of an effect were used to form clusters. For each cluster, we computed a cluster-level statistic by summing the t-values across the elements within the cluster. These clusters were then compared against a null distribution, which we generated from 10,000 sign-flip permutations. In each permutation, we flipped the sign of the accuracy scores for each subject, simulating a null effect. We repeated the same thresholding and clustering process for each permutation and recorded the maximum cluster statistic. This resulted in a null distribution representing the strongest clusters expected by chance. Each empirical cluster was then compared against this null distribution. Cluster-level p-values were computed as proportion of permuted maximum cluster statistics that were equal or exceed the empirical cluster statistic. Clusters with *p <* 0.05 were considered statistically significant.

## 3 Results

### 3.1 Behavioural Results

Behavioural results showed that our participants were engaging in the tasks in both Sessions. During Session 1 (Figure 2b), participants’ working memory performance was consistently high (*M* = 0.86, 95% CI [0.84, 0.88]). Performance decreased with difficulty level, from Easy (*M* = 0.94, 95% CI [0.92, 0.95]), to Medium (*M* = 0.86, CI [0.83, 0.89]), and to Hard (*M* = 0.78, CI [0.75, 0.80]). During Session 2, performance on the attentional control task was high (*M* = 0.98, 95% CI [0.96, 1]). For full details of all behavioural results see Supplementary Results (Appendix B.1).

### 3.2 FMRI Decoding Results

We first trained and tested classifiers on brain signals during perception (Session 2) (see Figure 3a). Classification accuracy for the images was significantly above chance (22.1%; 95% CI [19.8%, 24.5%], where chance level is 1.6%; *p <* 0.001). This corroborates that object-specific information can be reliably decoded from early visual cortex using our naturalistic experimental stimuli.

**Figure 3:**
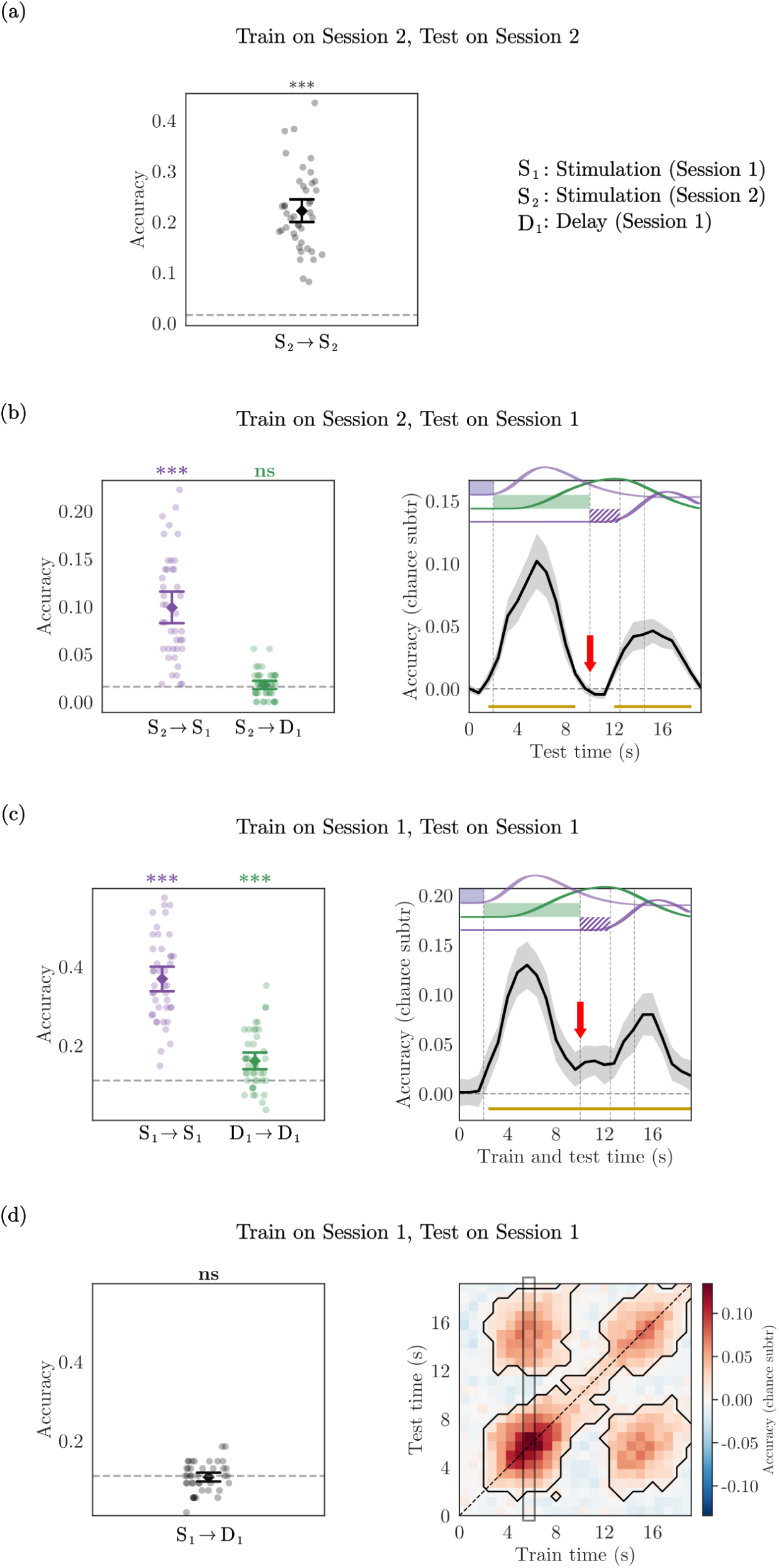
Within- and between-session decoding performance in early visual cortex. Arrows indicate the direction of decoding (*data_train_*→ *data_test_*). **(a)** GLM-based classification within Session 2. Cross-validated accuracy for sensory classification on GLM-parameter estimates trained and tested within Session 2 (*S*_2_ → *S*_2_). This shows robust above-chance decoding of sensory stimuli (\*\*\**p <* 0.001). The dashed horizontal line marks chance level (63 classes, ≈1.6%). **(b)** Between-session classification accuracy. Left: GLM-based decoding. A classifier was trained on Session 2 and applied to either the stimulation (*S*_2_ → *S*_1_) or delay period (*S*_2_ → *D*_1_) of Session 1. Classification of the stimulation period (purple) is robust, whereas there was no evidence for information for the delay period (green). Right: Time-based decoding. Classifiers were trained on data from Session 2 and tested on data from Session 1 for 20 s (25 TRs) following stimulus presentation onset (0 s). The filled purple, filled green and striped purple rectangles show the stimulation, delay and probe periods respectively. Thick black curves show the mean accuracy averaged across participants (grey bands: 95% CIs). Dark yellow bars along the x-axis highlight periods where classification accuracy significantly exceeded chance (one-sided sign-flip cluster-mass permutation test with 10,000 permutations and a cluster-defining threshold *p <* 0.05). Note the absence of decodable information based on the classifier trained on Session 2 towards the end of the delay period before onset of the probe (red arrow). For reference, the expected BOLD-response time courses for each condition are inserted at the top (based on an SPM canonical HRF, see Methods). The time course of the available information provides a good match to the expected sensory responses, but does not extend throughout the delay period and into the subsequent probe phase as would be expected from delay-period information. Note that this drop of information in the delay period only holds for the classifier that is trained on the purely sensory task in Session 2 (see subpanel c). **(c)** Within-session classification accuracy. Left: GLM-based decoding. A classifier was trained on either the stimulation or delay period of the Session 1 and applied to the same period (*S*_1_ → *S*_1_ and *D*_1_ → *D*_1_, respectively). Classification of the stimulation period (purple) and of the delay period (green) is above chance. Right: Time-based decoding. Mean classification accuracy over time with 95% confidence intervals, obtained when classifiers were trained and tested at the same timepoint. Chance level was subtracted before averaging across 42 participants. The dashed horizontal line indicates chance-level classification. Significant values from the cluster-mass permutation test (see panel b and Methods) are indicated by dark yellow bars. Horizontal rectangles and red arrow indicate task periods as in panel (b). **(d)** Within-session classification accuracy. Left: GLM-based decoding. A classifier was trained on the stimulation and applied to the delay period within Session 1 (*S*_1_ → *D*_1_). Classification did not differ from chance (*p* = 0.76). Right: Temporal generalisation analysis. Classifiers were trained and tested on repeated trials from Session 1 across a 25-TR (20 s) time window following stimulus onset. Heat map shows group-mean classification accuracies. Significant clusters, outlined with black contours, were determined using a maximum cluster-mass sign-flip permutation procedure. The grey vertical rectangle highlights the training timepoint yielding the highest decoding accuracy which, after accounting for the haemodynamic lag, corresponds to stimulus presentation. Classifiers trained at this and adjacent timepoints do not generalise throughout the delay period, but do generalise to timepoints relative to probe presentation (around 12 s).

We then proceeded to assess whether this percept-based classifier from Session 2 generalised to the decoding of stimulus identity in Session 1 (Figure 3b, left, purple). We first used a classifier trained on GLM parameter estimates from Session 2 and tested whether the classifier was able to decode stimulus identity in the stimulus-based period early in the trial in Session 1. Decoding accuracy was above chance (9.9%, 95% CI [8.2%, 11.5%], chance = 1.6%; *p <* 0.001), suggesting that the way stimulus information was encoded during the stimulation phase of the memory task (Session 1) was similar to the way it was encoded during the purely perceptual task (Session 2). Using the same percept-based classifier, we then tested the generalisation to the delay period (Figure 3b, left, green). In this case, classification accuracy was not significantly different from chance (1.8%, 95% CI [1.3%, 2.2%], chance = 1.6%; *p* = 0.224). Taken together these two results suggest that information from the perception-only condition generalises well to the stimulation phase but not to the delay phase of the working memory task.

In order to assess the temporal dynamics of this information in more detail, we conducted a time-resolved classification analysis, again testing for generalisation across sessions (Figure 3b, right). Classifiers were trained on raw BOLD signals from the perception-only session and tested at each timepoint of the memory-task trials. We found that classifiers generalised robustly during the early period of the trial, but information decayed very rapidly thereafter, reaching baseline during the delay about 9 seconds after stimulus onset. Several seconds after the presentation of the probes, stimulus decoding accuracy reached a second, albeit smaller peak. This presumably reflects the presentation of the probes which are highly similar to the targets. We also computed the expected latency periods when information should be present for each of the trial phases by accounting for the haemodynamic lag. The expected time courses are depicted in the top of Figure 3b (right). The first informative peak coincides with the expected haemodynamic delay of the stimulation phase, the second informative peak coincides with the expected haemodynamic delay of the probe phase. In contrast, the gap where decoding accuracy fell back to baseline coincided with the expected peak of potential memory signal, suggesting that the sensory-based classifier was not informative about the representations maintained across the entire delay period.

One possibility is that the low classifier performance in the later delay period reflects the overall weaker information of memory as compared to sensory signals (Duan & Curtis, 2024). Alternatively, the information might continue to be encoded, albeit in a format that is different from the sensory format in Session 2 (and the stimulation phase of Session 1). To assess this we first trained classifiers using GLM parameter estimates from Session 1 (Figure 3c, left) and applied them to stimulation and delay phases separately. Classification in the stimulation phase was above chance (36.9%, CI [33.6%, 40.1%], chance = 11%; *p <* 0.001), in line with our previous analyses. In contrast to the percept-based classifier, however, classification from the delay phase was also significantly above chance (16.1%, CI [13.9%, 18.3%], chance = 11%; *p <* 0.001). This suggests that working memory contents can indeed be decoded from the delay phase, only that they are not in a sensory format. In order to assess the temporal dynamics of this information, we again conducted a time-resolved analysis, but now only within Session 1. Classifiers were trained on raw BOLD signals at each timepoint separately over the course of a trial and were tested on matching timepoints (Figure 3c right). In line with our findings from the GLM model (Figure 3c, left) classifiers were able to decode stimulus-specific information throughout the trial, including throughout the delay phase. We conducted a supplementary whole-brain searchlight analysis based on GLM parameter estimates to identify regions carrying stimulus information during the delay period (Appendix B.4, Figure S14). The analysis revealed significant clusters in early visual cortex, providing converging evidence this region is engaged in working memory maintenance.

If the stimulus presentation and delay phases of the working memory task indeed encode information in a different format, cross-decoding from stimulus presentation to delay phase should be substantially reduced or even at chance. We thus conducted an additional analysis, again on GLM parameter estimates from Session 1 (Figure 3d, left), training on the stimulation period and testing on the delay period. Decoding accuracy was at chance (10.8%, CI [9.6%, 11.9%], chance = 11%; *p* = 0.76), confirming that the perception-based classifiers failed to generalise to the delay-phase independently of whether classification was within- or between-sessions. Finally, we conducted a further temporal generalisation analysis on raw BOLD data from Session 1, to obtain a comprehensive characterisation of how classifiers trained at each timepoint within a trial generalise across all other timepoints (Figure 3d, right). Within-time classification accuracies (i.e. the diagonal where training and testing are done on the same timepoint) rose above chance following stimulus onset, peaked between 4.8 and 6.4 s, and then declined but remained above chance for the remainder of the trial. A modest secondary peak emerged during the delay period, followed by a more pronounced increase after probe presentation. As in our previous analyses we did not find any evidence for cross-temporal generalisation from encoding into the delay period (cf. Figure 3d, right, vertical grey rectangle), whereas cross-temporal generalisation between the stimulus and probe presentation was robust in both directions. Note that the presence of perceptually decodable memory information early in the trial cannot be ruled out (see Discussion). These results were replicated using a correlation-based decoder instead of SVMs, as well as across all occipital and parietal ROIs in a subsequent exploratory analysis (see Supplementary Results, Appendix B.2, and B.3).

## 4 Discussion

In this study, we compared how early visual cortex encodes stimulus-specific information during perception and a working memory delay. We recorded participants’ brain signals using fMRI over two consecutive sessions. In the first session, they were asked to memorise naturalistic object-scenes over a delay, whereas in the second session they performed a purely perceptual task. As expected, we found that the stimuli could be decoded from early visual cortex in the perceptual task. The same sensory-based classifier trained on Session 2 was able to reliably decode information throughout the early phase of the working memory task in Session 1, but this did not extend throughout the entire delay. When instead using classifiers trained on the matching time periods of the memory task itself the stimulus was decodable throughout in Session 1. This suggests that the information in the early phase of the working memory experiment was perception-like, but this did not last throughout the delay. Thus, the way information was encoded in the later phase of the delay period differed from a pure sensory code. Our findings indicate that working-memory representations are not strictly sensory-like, and provide an important refinement of sensory recruitment models of visual working memory.

A key feature of our experimental design was that we recorded brain activity during a purely perceptual task in a separate session (see our discussion of related studies below; Harrison & Tong, 2009; Albers et al., 2013; Rademaker et al., 2019; Iamshchinina et al., 2021). This enabled us to separate between sensory-like and other types of representations using an independent perceptual training set. The sensory information decoded from the early phases of the working memory task presumably reflects the initial sensory processing of the memorised item. This is confirmed by two complementary analyses where classifiers were trained on the perceptual task in Session 2 and then applied to the working memory task in Session 1. The GLM-based analysis only identified sensory information for the stimulus presentation, but not for the delay (where it was at chance level). A more detailed time-resolved classifier showed two clear peaks which, accounting for the HRF lag, correspond to stimulus presentation and probe presentation. The latter is lower in magnitude, likely due to the fact that in half of the trials the first of the two probes is the foil. Between the sensory and the probe-related peaks, the classification accuracy rapidly goes back to baseline, suggesting that the early visual cortex does not maintain information in the same format throughout the delay phase. If this were the case, classification accuracy should have remained above chance between the two peaks. The lack of cross-decodability cannot be explained by a lack of sensitivity, as the classifiers generalised robustly across sessions during the other periods. The high accuracy during these other periods indicates that our naturalistic object-scene stimuli evoked robust and easily decodable sensory representations in the early visual cortex despite the large stimulus set. Taken together, the GLM-based and the temporal generalisation analysis yielded converging results.

Please note that our results do not suggest that there is no decodable information about the stimuli in early visual cortex during working memory maintenance. Quite the contrary, we could reliably decode object identity from the working memory delay, but only when classifiers trained on the matching delay period were used (both using GLM parameter estimates and time-resolved). Information in the later delay stages was lower than in the earlier stages. It has long been known that working memory representations are not full-blown sensory representations and most studies report a drop in decoding accuracy throughout the delay (Christophel et al., 2012; Li & Curtis, 2023; Degutis et al., 2025). Especially the time-resolved analyses showed that stimulus decodability remained above chance for the whole duration of the trial when the same (or neighbouring, see Figure 3d) timepoints were used to train and test the classifiers. We interpret this as evidence that the early visual cortex carries stimulus-specific information throughout the working memory delay, but that this format changes over time, departing from a sensory-like code. Importantly, we cannot exclude that memory-related information in the early period of the working memory delays is sensory-like. The information we found in the early phase could either reflect purely the sensory responses to the target stimuli at the beginning of the trials. Alternatively, the code in which working memory contents are encoded might exhibit non-stationarity across the delay (e.g. Sreenivasan et al. (2014)), being more sensory-like in the early phase and differing from sensory encoding in the later stage. Thus, our results are compatible with recent emphasis on dynamic coding in the early visual cortex (Meyers et al., 2008; Stokes et al., 2013; Sreenivasan et al., 2014; Spaak et al., 2017; Meyers, 2018; Li & Curtis, 2023; Degutis et al., 2025), which support the idea of working memory representations being reformatted during maintenance.

Our results do not fundamentally challenge the sensory recruitment model of visual working memory (Awh & Jonides, 2001; Postle, 2006; D’Esposito, 2007; Serences et al., 2009), they just point to specific clarifications. This is because early visual cortex still has substantial stimulus-related information throughout the working memory delay. What our findings suggest is that the encoding of this information is not necessarily sensory-like. To put it simply: sensory areas still seem to be recruited for visual working memory, but their recruitment might occur in a more flexible fashion using different, non-sensory encoding schemes. Our study is not the first to assess how percept-based classifiers generalise to working memory delays.

Several previous studies have found significant cross-decodability between sensory and working memory tasks (e.g. Harrison and Tong (2009), Albers et al. (2013), Rademaker et al. (2019), Iamshchinina et al. (2021), and Vo et al. (2022)). Typically, the information also decays throughout the delay period (Harrison & Tong, 2009; Albers et al., 2013; Rademaker et al., 2019; Iamshchinina et al., 2021). What our study suggests is that the perception-like information can be completely absent at the end of the delay, raising the question how sensory contents are maintained until the retrieval. The notion that memory maintenance relies on a sustained sensory code has recently faced scrutiny also in other studies (Duan & Curtis, 2024). In their study, Duan and Curtis (2024) designed an experimental manipulation with a specific variant of gratings to compare the cross-decodability of sensory and memory codes, while accounting for vignetting effects of grating stimuli (Roth et al., 2018). While sensory representations in visual cortex were specific to each modulated grating, working memory representations generalised across them, which corroborates the notion of working memory codes being an abstraction of sensory ones (Kwak & Curtis, 2022). Another recent study by Xu (2023) using grayscale object stimuli found a significant drop in categorical decoding between stimulation and delay phases across occipital and parietal areas, suggesting that, for real-world objects, sensory and mnemonic content might have different representational geometries. Our results suggest that working memory codes are not only a “weaker” version of sensory ones, but rather that they undergo significant reformatting. Although our design did not allow us to explicitly test whether the reformatting of working memory representations corresponds to an abstraction of sensory codes as in Duan and Curtis (2024), our results point in the same direction.

A key difference between our study and the abovementioned work, is that we used naturalistic images instead of orientation gratings or other low-dimensional stimuli. One could therefore wonder, whether the lack of sensory-to-memory generalisation is explained by our stimuli being semantically rich, and their encoding representations being consequently maintained in higher order visual areas. Two results speak against this hypothesis. First, our design ensured that participants could only successfully perform the working memory task by relying on fine-grained perceptual discrimination. Our stimulus design ensured that target and foils were only gradually different in terms of fine-grained details while maintaining overall semantic information (see Figure 2). Thus, only memorizing the overall category or even the “gist” of the images (Friedman, 1979) would not be sufficient to identify the correct target. Instead, the fine-grained spatial details encoded in early visual areas would provide very useful information in solving the task. A second reason why working memory for our naturalistic stimulus task does not appear to be delegated to further downstream areas is provided by a supplementary ROI comparison (Supplementary Results, Appendix B.2, Figure S12). The early visual cortex yielded higher accuracies for both sensory and working memory classification than downstream object- and category-selective regions, which further confirms that our task primarily engaged perceptual discrimination rather than semantic processing.

It has been proposed that the role of visual cortex in working memory maintenance could depend on task demands (Serences, 2016). According to this hypothesis (C. S. Adam et al., 2022), if the task requires participants to store sensory features, the visual cortex will rely on sensory codes. On the other hand, if the task demands are more abstract, abstract codes will be used. This hypothesis is based on the idea that the visual cortex can switch between different sensory and mnemonic codes (Rademaker et al., 2019), and supports a conception of working memory as a distributed function (Christophel et al., 2017). The fact that object-identity was best decoded from the early visual cortex in our study supports the idea that our task-demands, which required focusing on fine-grained details in the stimuli, involved representations in early visual cortex. Experimental tasks with category-based demands (other than in our study), for example, tend to find more stimulus-specific working memory information and some sensory-memory generalisation in category-selective areas (e.g. lateral occipito-temporal and ventral occipito-temporal cortex) than in the early visual cortex (e.g. Xu (2023)). However, our results do point to a refinement of the sensory recruitment model. While our task demanded the storage of fine-grained perceptual information across the delay phase, and while the early visual cortex carried robust stimulus-specific information throughout the delay phase, the way in which the stimulus was represented appeared to change. The fact that sensory-trained classifiers did not generalise throughout the delay period suggests a representational discontinuity between perception and working memory maintenance. This could potentially be explained by the necessity to shield the memory representation from incoming perceptual interference. Our results are more compatible with an orthogonalisation of sensory and memory codes (Libby & Buschman, 2021) rather than a parallel coexistence in the same format (Rademaker et al., 2019).

Our approach using naturalistic images, moreover, fills a gap in the literature. Previous studies have mostly used simple stimuli with minimal semantic meaning (but see Lee et al. (2012), Lee and Kuhl (2016), and Xu (2023)), raising a problem of generalisability of sensory recruitment to real-world environments, which are perceptually rich and semantically meaningful. Despite a growing interest in a more naturalistic understanding of working memory processes (Brady et al., 2019; Xu, 2023; Bates et al., 2024; Adam et al., 2025), there is a lack of evidence for sensory recruitment using naturalistic images. Thus, our findings extend the existing literature by showing that the early visual cortex contains a memory signal also for perceptually rich, naturalistic images. A common experimental limitation when using naturalistic stimuli for working memory tasks is calibrating target-foil similarity to threshold-level to avoid ceiling and floor effects. We tackled this challenge by using a synthetically generated stimulus set (Pettini et al., 2025), which allowed us to parametrically vary target-foil perceptual similarity while keeping semantic identity constant (e.g. variations of “a picture of a cat on the carpet”). Our task required the participants to reidentify the target among other foils of similar spatial layout and semantic meaning. Thus, a verbal or category-based strategy would not have been helpful. The success of the experimental manipulation was confirmed both by behavioural and neuroimaging results. Behaviourally, results showed that participants’ accuracies decreased in a graded fashion as target-foil similarity increased. Our result is consistent with behavioural models linking memory signal strength and perceptual discriminability (Schurgin, 2020), extending them to naturalistic stimuli. An additional set of exploratory analyses in higher order visual regions revealed that, despite the stimuli being semantically meaningful, classification accuracy was strongest in the early visual cortex, corroborating the idea that participants attended to perceptual details. In naturalistic cognition, the visual system processes complex images, rich in perceptual and semantic detail. Laboratory settings aiming at precise experimental control have so far privileged low-dimensional, non-semantic stimuli. Our findings show that if one wants to exhaustively understand the role of the early visual cortex recruitment in working memory maintenance, complementary evidence using naturalistic stimuli is necessary.

One open question is the causal necessity of the early visual cortex for memory maintenance. Critics have highlighted a divergence between human fMRI and non-human primate electrophysiology, which rarely detect sustained, stimulus-specific activity in early sensory areas (Leavitt et al., 2017; Curtis & Sprague, 2021), thus indicating a potential segregation of sensory and memory circuitries (Roussy et al., 2021). Additionally, some argued that because the working memory signal observed in the early visual cortex can be driven by top-down modulation from higher-order areas (Lawrence et al., 2018) rather than local storage, the early visual cortex may not be functionally essential for memory maintenance (Bettencourt & Xu, 2016; Xu, 2020). However, the presence of feedback signals from higher order areas does not make early visual cortex signals epiphenomenal. The interaction can also be interpreted as an active recruitment mechanism, where higher order areas utilise early visual cortex circuitry as a high-resolution buffer (Roelfsema & de Lange, 2016; van Kerkoerle et al., 2017). Our study was not designed to address this question directly, and future work should elucidate the dynamics between early visual cortex and higher cortical areas during working memory maintenance. However, we did observe that early visual cortex had a maximum of information about the memorized stimulus when compared to other regions.

Here we show that, using naturalistic object-scenes and a task with perceptual demands, we find object-specific information in the early visual cortex, both during perception and working memory maintenance. However, the sensory-to-memory generalisation does not last throughout the delay phase, providing evidence for at least partial segregation of these representations. Our findings challenge the view that sensory recruitment entails a simple reinstatement of sensory-like codes, and supports the hypothesis that the early visual cortex maintains sensory working memory contents in reformatted, non-sensory representations.

## Declarations

### Ethics approval

The studies were approved by the Ethics Committee of the Institute of Psychology of the Humboldt University Berlin, Germany.

### Consent to participate

All participants provided informed written consent prior to participation in the study, in accordance with institutional guidelines.

### Consent for publication

Participants were informed that anonymised data may be published in scientific journals and provided consent for publication.

## Appendix A: Supplementary Methods

### A.1 Behavioural modelling

We analysed participants’ trial-level accuracy using a Bayesian hierarchical logistic regression model, implemented with the brms package (Bürkner, 2017) in R (R Core Team, 2024). The primary goal of this analysis was to evaluate whether participants’ performance was affected by task difficulty, and also whether it improved across runs. We also investigated if stimulus repetition improved working memory performance. The log-odds of a correct response were modelled as a function of run number (1 to 6), task difficulty (Easy, Medium, Hard) and target repetition (repeated vs non-repeated). We included every two-way interaction and the three-way interaction among these factors, as well as participant-specific intercepts and slopes to model, respectively, baseline performance and individual learning rates. The Bayesian model was estimated via Markov chain Monte Carlo (MCMC) sampling using four chains of 10,000 iterations each. The first 2,000 iterations per chain were discarded as warm-up. Across all chains, we ran a total of 32,000 post-warm-up samples. The model was fit for 4,472 trials. We assessed convergence using the potential scale reduction statistic (*R*^^^) (Gelman & Rubin, 1992), and we evaluated chain resolution using the effective sample sizes (ESS) for all parameters (Kruschke, 2021). We modelled the factor “run number” as a continuous variable from 1 to 6, “target condition” as a two-level factor (repeated versus non-repeated), and difficulty as a three-level factor (Easy, Medium, Hard). In the model, we included all two-way interactions (run number x difficulty, run number x target condition, difficulty x target condition) and the three-way interaction (block x difficulty x target condition).

For each trial *i* completed by participant *j*, we modelled the binary outcome variable *y_ij_*(indicating whether the response was correct) as a Bernoulli-distributed variable:

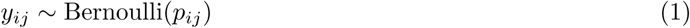

with the probability of a correct response, *p_ij_*, linked to a linear predictor using the logit function:

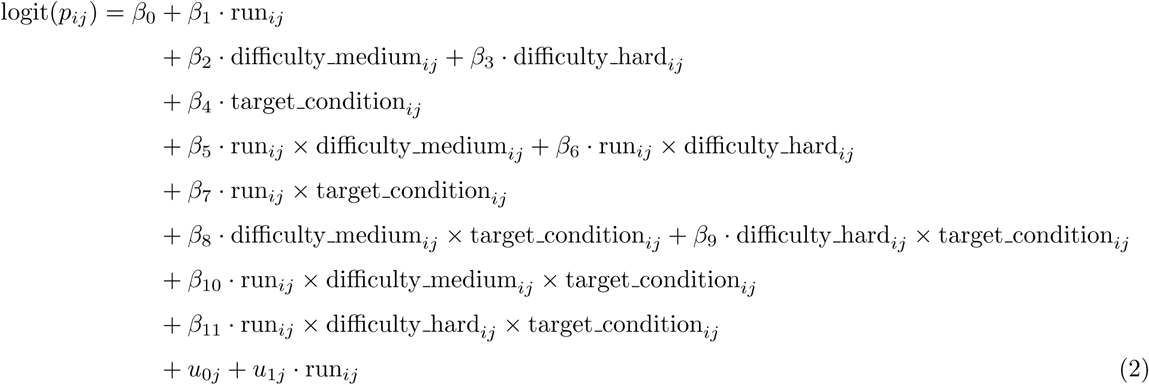

where *β*_0_ corresponds to the log-odds of a correct response under the reference condition (that is, first run, Easy difficulty, non-repeated targets). The other *β* coefficients quantify the change in log-odds relative to the baseline, estimating the fixed effects of run, difficulty, target condition, and their interactions. The random intercepts *u*_0_*_j_* and slopes *u*_1_*_j_* model between-participant variability in baseline performance and improvement throughout the task. They were allowed to correlate with covariance matrix:

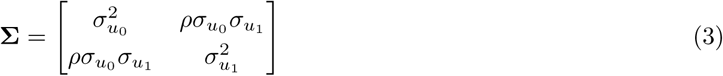

Here, *σ_u_*_0_ and *σ_u_*_1_ are the standard deviations of the random intercept and slope, whereas *ρ* represents the correlation between the two.

As in Pettini et al. (2025), we specified weakly informative priors to regularise parameter estimation and reduce the risk of overfitting (Gelman, 2006). For the fixed effects, we used normal priors:

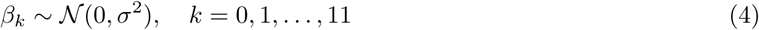

with *σ*^2^ = 4, which constrains the coefficients within a range of approximately ±4 at two standard deviations on the log-odds scale. For the standard deviations of the random effects (*σ_u_*_0_ and *σ_u_*_1_), we used half-Cauchy priors:

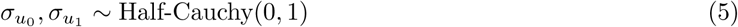

To model the correlation between random intercepts and slopes (*ρ*), we used a Lewandowski-Kurowicka-Joe (LKJ) prior:

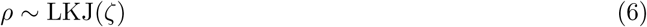

with shape parameter *ζ* = 2, which discourages extreme correlations unless supported by the data (Lewandowski et al., 2009; Bürkner, 2017).

We ran prior predictive checks to ensure the priors generated reasonable simulated outcomes. To assess robustness, we also analysed alternative weakly informative priors. Posterior predictive checks were used to evaluate the model fit.

### A.2 Preprocessing (fmriprep boilerplate)

#### Copyright Waiver

The above boilerplate text was automatically generated by fMRIPrep with the express intention that users should copy and paste this text into their manuscripts *unchanged*. It is released under the CC0 license.

Results included in this manuscript come from preprocessing performed using *fMRIPrep* 24.1.0rc0 (Esteban, Markiewicz, Blair, Moodie, Isik, Erramuzpe Aliaga, et al. (2019); Esteban et al. (2018); RRID:SCR 016216), which is based on *Nipype* 1.8.6 ((Gorgolewski et al., 2011); (Gorgolewski et al., 2018); RRID:SCR 002502).

#### Anatomical data preprocessing

A total of 4 T1-weighted (T1w) images were found within the input BIDS dataset. Each T1w image was corrected for intensity non-uniformity (INU) with N4BiasFieldCorrection (Tustison et al., 2010), distributed with ANTs 2.5.3 (Avants et al., 2008, RRID:SCR 004757). The T1w reference was then skull-stripped with a *Nipype* implementation of the antsBrainExtraction.sh workflow (from ANTs), using OASIS30ANTs as the target template. Brain tissue segmentation of cerebrospinal fluid (CSF), white matter (WM), and grey matter (GM) was performed on the brain-extracted T1w using fast (FSL (version unknown), RRID:SCR 002823, Zhang et al., 2001). An anatomical T1w reference map was computed after registration of the 4 images (after INU correction) using mri robust template (FreeSurfer 7.3.2, Reuter et al., 2010). Volume-based spatial normalization to two standard spaces (MNI152NLin6Asym, MNI152NLin2009cAsym) was performed through nonlinear registration with antsRegistration (ANTs 2.5.3), using brain-extracted versions of both the T1w reference and the T1w template. The following templates were selected for spatial normalization and accessed with *TemplateFlow* (24.2.0, Ciric et al., 2022): *FSL’s MNI ICBM 152 non-linear 6th Generation Asymmetric Average Brain Stereotaxic Registration Model* [(Evans et al., 2012), RRID:SCR 002823; TemplateFlow ID: MNI152NLin6Asym], and the *ICBM 152 Non-linear Asymmetrical template version 2009c* [(Fonov et al., 2009), RRID:SCR 008796; TemplateFlow ID: MNI152NLin2009cAsym].

#### Functional data preprocessing

For each of the 14 BOLD runs found per subject (across all tasks and sessions), the following preprocessing was performed. First, a reference volume was generated, using a custom methodology of *fMRIPrep*, for use in head motion correction. Head-motion parameters with respect to the BOLD reference (transformation matrices and six corresponding rotation and translation parameters) were estimated before any spatiotemporal filtering using mcflirt (FSL Jenkinson et al., 2002). The BOLD reference was then co-registered to the T1w reference using mri coreg (FreeSurfer) followed by flirt (FSL Jenkinson & Smith, 2001) with a boundary-based registration (Greve & Fischl, 2009) cost function. Co-registration was configured with six degrees of freedom. Several confounding time series were calculated based on the *preprocessed BOLD*: framewise displacement (FD), DVARS, and three region-wise global signals. FD was computed using two formulations: Power (absolute sum of relative motions, (Power et al., 2014)) and Jenkinson (relative root mean square displacement between affines, (Jenkinson et al., 2002)). FD and DVARS were calculated for each functional run, both using their implementations in *Nipype* (following the definitions by Power et al., 2014). Global signals were extracted within the CSF, white matter, and whole-brain masks. Physiological regressors were extracted to allow for component-based noise correction (*CompCor*, Behzadi et al., 2007b). Principal components were estimated after high-pass filtering the *preprocessed BOLD* time series (using a discrete cosine filter with a 128 s cut-off) for the two *CompCor* variants: temporal (tCompCor) and anatomical (aCompCor). tCompCor components were calculated from the top 2% most variable voxels within the brain mask. For aCompCor, three probabilistic masks (CSF, working memory, and combined CSF+working memory) were generated in anatomical space. The implementation differs from that of Behzadi et al. in that instead of eroding the masks by 2 pixels in functional space, a mask of voxels including a small fraction of GM was subtracted, obtained by thresholding the partial volume map at 0.05. Masks were then resampled into BOLD space and binarized at 0.99 (as in the original implementation). Components were calculated separately within working memory and CSF masks. For each CompCor decomposition, the *k* components with the largest singular values were retained such that they explained 50 percent of the variance across the nuisance mask. Head-motion estimates from the correction step were also stored in the corresponding confounds file. Confound time series derived from motion estimates and global signals were expanded with temporal derivatives and quadratic terms (Satterthwaite et al., 2013). Frames exceeding 0.5 mm FD or 1.5 standardised DVARS were annotated as motion outliers. Additional nuisance time series were calculated using PCA on a thin band (*crown*) of voxels around the brain edge (Patriat et al., 2017). All resampling operations were performed with *a single interpolation step* by composing all relevant transformations (head-motion matrices, susceptibility distortion correction when available, and co-registrations to anatomical and output spaces). Volumetric resampling was performed using nitransforms, configured with cubic B-spline interpolation.

Many internal operations of *fMRIPrep* use *Nilearn* 0.10.4 (Abraham et al., 2014, RRID:SCR 001362), mostly within the functional processing workflow. For more details of the pipeline, see the section corresponding to workflows in *fMRIPrep*’s documentation.

### A.3 Correlation-based Decoder

Sensory and mnemonic patterns may share their multivariate geometry despite having different state-dependent baseline shifts. To make our analysis sensitive to these geometry-preserving correspondences, which standard linear classifiers might fail to capture, we implemented a time-resolved correlation-based decoder. Raw voxel time-series were extracted per ROI and preprocessed using the same method described above (run-wise linear detrend and voxel-wise z-normalisation). For Session 2, we derived one mean activation pattern per label (object-scene) by averaging its 8 activation patterns (one presentation per run, 8 runs). For Session 1, we extracted a 25-TR (20 s) window per trial time-locked to stimulation onset. For every trial, we computed the Pearson’s correlation coefficient between the multivoxel pattern at every timepoint and each of the 63 Session-2 mean activation patterns. We did this for all trials (repeated and non-repeated). In case of repeated trials, we calculated the correlation coefficients separately for each repetition, applied the Fisher z-transformation, averaged and then inverse-transformed. For each trial and timepoint, we thus obtained one correlation value per label (63 in total). To calculate classifier accuracy, we extracted the label with the highest correlation value at each TR, compared it to the true label of the current trial, and classified matches as correct and mismatches as incorrect. Finally, we computed the mean accuracy and 95% confidence intervals at each time-point across participants. Statistical significance was assessed using the same non-parametric procedure described in (Section 3.6.6).

### A.4 Searchlight Analyses

We ran a set of exploratory whole-brain searchlight analyses (Kriegeskorte et al., 2006) to identify clusters carrying stimulus-specific information beyond the ROIs. The analysis was run using single-trial beta maps as input features and a leave-one-run-out cross-validation scheme. We used spherical searchlights of radius 4 mm defined in subject space and included in a brain mask. For each participant, we obtained a classification accuracy map in which the value at each voxel reflects the classification accuracy achieved when using a sphere centred on that voxel. Chance level was subtracted voxel-wise from the resulting accuracy maps, which were then saved and transformed to MNI space for group analysis, which was performed using SPM12 (https://www.fil.ion.ucl.ac.uk/spm/software/spm12/). To assess statistical significance, we performed a second-level one-sample t-test across subjects to test whether classification accuracy at each voxel was significantly above chance. Multiple comparisons were corrected using FWE cluster correction at *p <* 0.05, with a voxel threshold of *p <* 0.001.

### A.5 Regions of Interest

**Figure S1:**
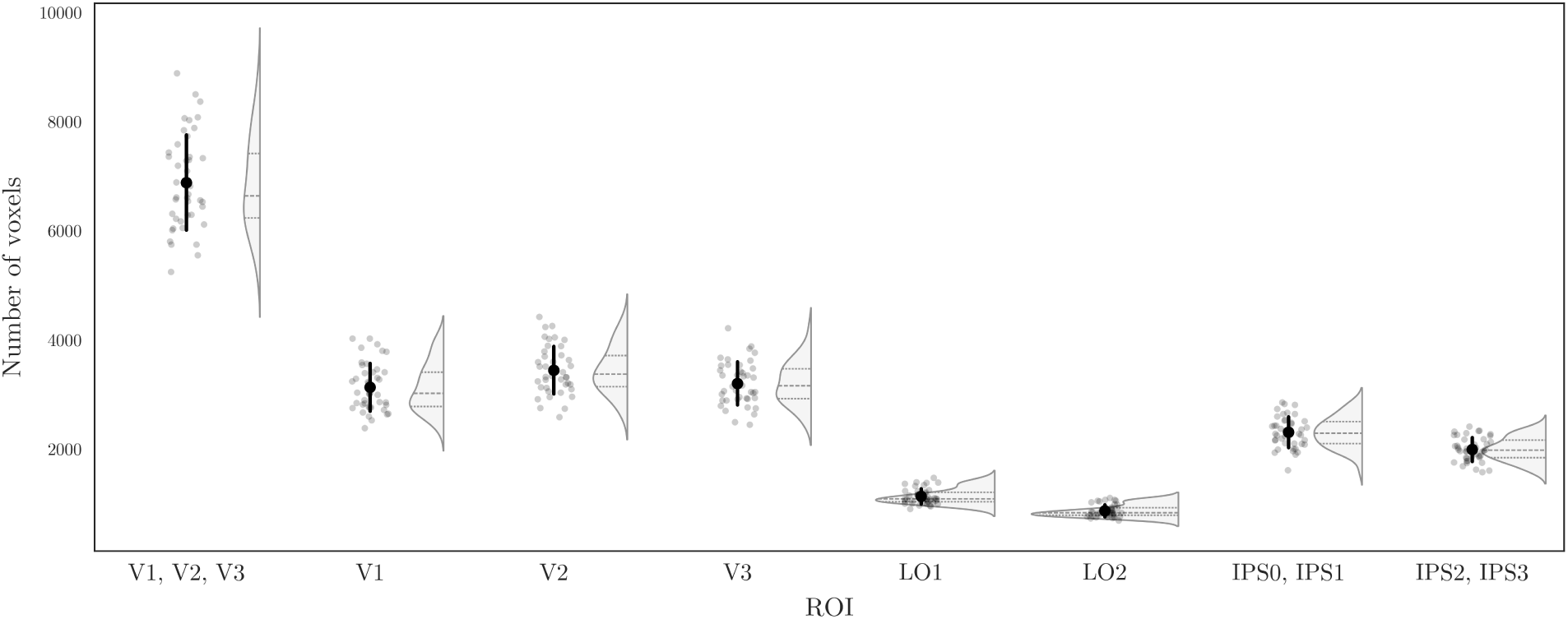
Voxel counts per visual ROI. Violin plots display the distribution of voxel counts across participants (gray dots). Black dots and error bars represent mean ± standard deviation. Visual ROI masks from Wang et al. (2015) were warped to native space, thresholded at 10%. Voxel counts were computed in the T1-aligned functional reference space (2-mm isotropic resolution).

## Appendix B: Supplementary Results

### B.1 Behavioural Results

We fitted a Bayesian hierarchical logistic regression predicting trial-wise accuracy from run number, task difficulty (Easy, Medium, Hard) and target repetition (Novel vs. Repeated), with by-participant random intercepts and slopes for run number. The model was estimated on 4,472 trials from 42 participants using four chains of 10,000 iterations each (2,000 warm-up; 32,000 post-warm-up draws). The model fitting showed robust convergence (*R*^^^ = 1.00, Bulk ESS *>* 9000, Tail ESS *>* 16000) (see Table 1). Participants’ behavioural accuracy was higher when targets and foils were more discriminable: OR = 0.22, 95% CI [0.11, 0.42] for Hard versus Easy, and OR = 0.44, 95% CI [0.22, 0.87] for Medium versus Easy (confidence intervals exponentiated from log-odds). The main practice effect across runs was small and uncertain, OR = 1.06, 95% CI [0.90, 1.23]. By contrast, we found strong evidence that repetition increased learning across runs (*β* = 0.28, 95% CI [0.07, 0.50], ER = 70.91, *pp* = .99), corresponding to a 32% greater increase in odds of a correct response per run. No other two- or three-way interactions showed credible deviations from zero. Participants varied moderately in their starting accuracy (SD = 0.46, 95% CI [0.21, 0.73]) but showed less variation in their improvement across runs (SD = 0.09, 95% CI [0.01, 0.18]). We did not find any systematic relationship between initial accuracy and improvement over runs (correlation = −0.11, 95% CI [−0.71, 0.71]). Baseline performance on Easy, non-repeated trials was high (*β* = 2.34, 95% CI [1.77, 2.94]), corresponding to a probability of success of approximately .91 (OR = 10.4). Trial difficulty produced a progressive reduction in performance: relative to Easy trials, Hard trials showed a 78% decrease in odds of a correct response (*β* = −1.50, 95% CI [−2.17, −0.85]; OR = 0.22), and Medium trials a 56% decrease (*β* = −0.81, 95% CI [−1.52, −0.13]; OR = 0.44). Repetition had no credible main effect on its own (*β* = −0.22, 95% CI [−1.06, 0.63]; OR = 0.80), but the interaction between run number and repetition was credibly positive (*β* = 0.28, 95% CI [0.03, 0.54]; OR = 1.32). This result, indicating that repeated targets gained a further 32% increase in odds of a correct response per run compared with novel targets, suggests that participants’ accuracy benefited from target repetition. All other two- and three-way interactions had 95% CIs spanning zero, providing no evidence of additional modulation by difficulty.

**Figure S2:**
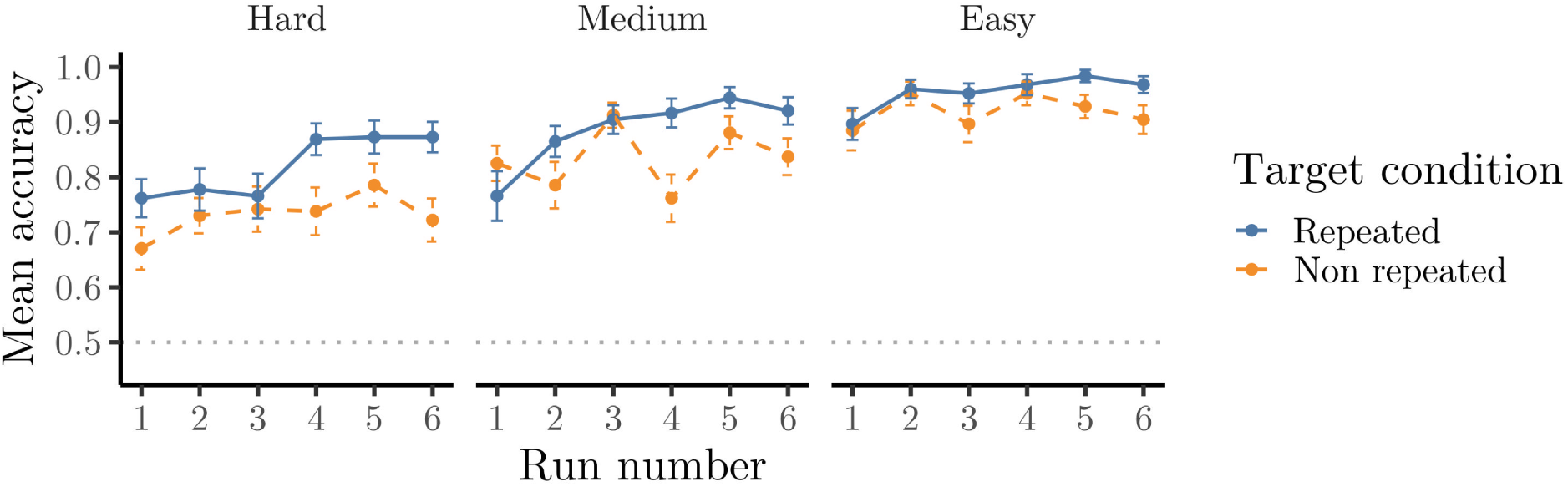
Behavioural performance across runs and difficulty levels. Mean recognition accuracy over runs grouped by difficulty condition (Hard, Medium, Easy). Bars are 95% CIs.

**Figure S3:**
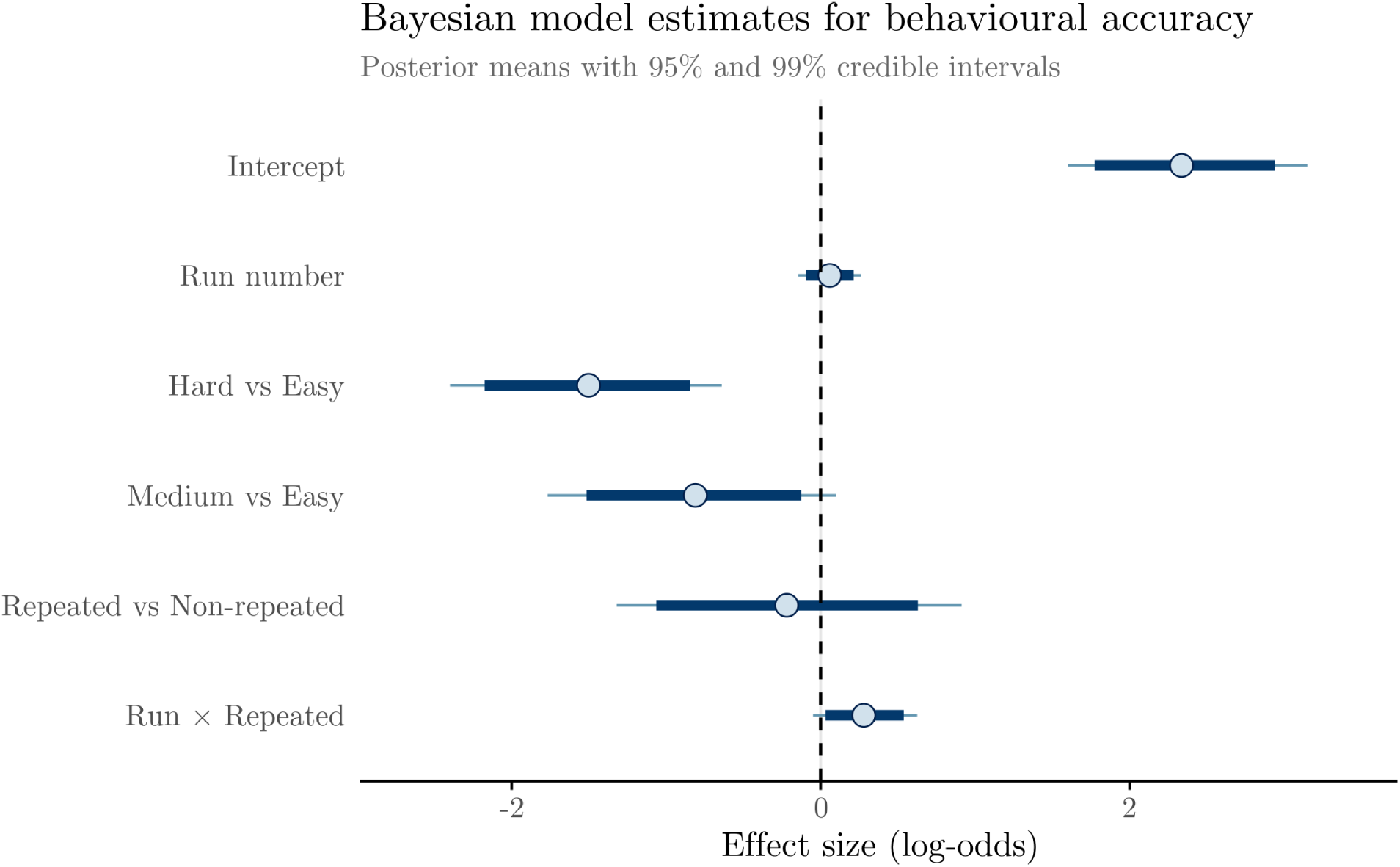
Bayesian hierarchical logistic-regression estimates for behavioural accuracy. Points depict the posterior means, thick and thin bars represent respectively 95% and 99% credible intervals. Dashed line represents no effect. Hard and Medium trials show progressively lower accuracy relative to Easy trials. Repetition shows no credible main effect, but the Run × Repeated interaction is positive, indicating greater improvement for repeated than for non-repeated targets across runs.

**Figure S4:**
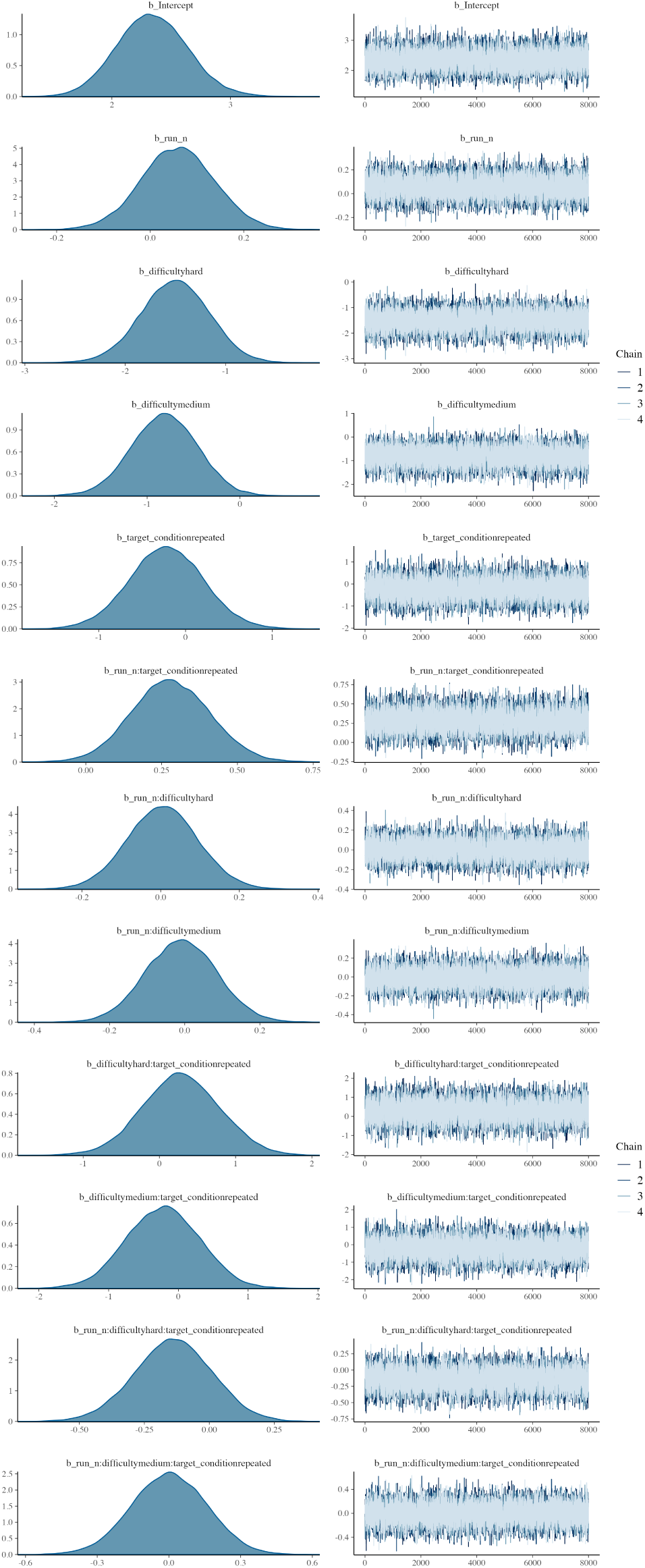
Posterior densities (left) and MCMC trace plots (right) for the Bayesian hierarchical logistic regression model.

**Table 1:** Posterior Summary of Fixed Effects.

| Parameter | Estimate | Est. Error | 95% CI Lower | 95% CI Upper | Rhat | Bulk ESS | Tail ESS |
| --- | --- | --- | --- | --- | --- | --- | --- |
| Intercept | 2.342 | 0.298 | 1.774 | 2.941 | 1.000 | 10799.628 | 17984.539 |
| run_n (Learning Effect) | 0.059 | 0.078 | -0.095 | 0.214 | 1.000 | 10292.362 | 17994.837 |
| Hard Difficulty (vs. Easy) | -1.504 | 0.339 | -2.175 | -0.847 | 1.000 | 11007.549 | 17881.175 |
| Medium Difficulty (vs. Easy) | -0.813 | 0.356 | -1.516 | -0.125 | 1.000 | 10909.179 | 18827.890 |
| Repeated Condition (vs. Non-Repeated) | -0.218 | 0.429 | -1.064 | 0.629 | 1.000 | 9286.196 | 16850.391 |
| run_n × Hard Difficulty | 0.005 | 0.090 | -0.171 | 0.183 | 1.000 | 10493.578 | 18505.494 |
| run_n × Medium Difficulty | -0.009 | 0.094 | -0.193 | 0.176 | 1.000 | 10633.438 | 18238.061 |
| run_n × Repeated Condition | 0.281 | 0.129 | 0.032 | 0.538 | 1.000 | 9497.827 | 16786.217 |
| Hard Difficulty × Repeated Condition | 0.286 | 0.506 | -0.705 | 1.281 | 1.000 | 9652.255 | 16913.457 |
| Medium Difficulty × Repeated Condition | -0.219 | 0.525 | -1.246 | 0.795 | 1.000 | 9908.217 | 17075.068 |
| run_n × Hard Difficulty × Repeated Condition | -0.139 | 0.149 | -0.435 | 0.151 | 1.000 | 9690.640 | 16922.793 |
| run_n × Medium Difficulty × Repeated Condition | 0.002 | 0.157 | -0.308 | 0.306 | 1.000 | 10216.147 | 17646.249 |

We also ran directed Bayesian hypothesis tests using Evidence Ratios (ERs) and corroborated these findings (Table 2). We found strong support for decreased performance on Hard versus Easy trials (H2: ER = ∞, *pp* = 1.00) and on Medium versus Easy trials (H3: ER = 90.43, *pp* = 0.99). There was weak evidence for a positive effect of run number (H1: ER = 3.44, *pp* = 0.77) and no evidence for a main effect of repetition (H4: ER = 0.44, *pp* = 0.30). In contrast, there was strong support for the run-by-repetition interaction (H5: ER = 70.91, *pp* = 0.99). All other interaction hypotheses involving difficulty received little to no support (ERs ≤ 2.52; *pp* ≤ 0.72). These results confirm robust graded difficulty effects and a selective practice-related gain for repeated targets, with no credible evidence for higher-order interactions.

**Table 2:**
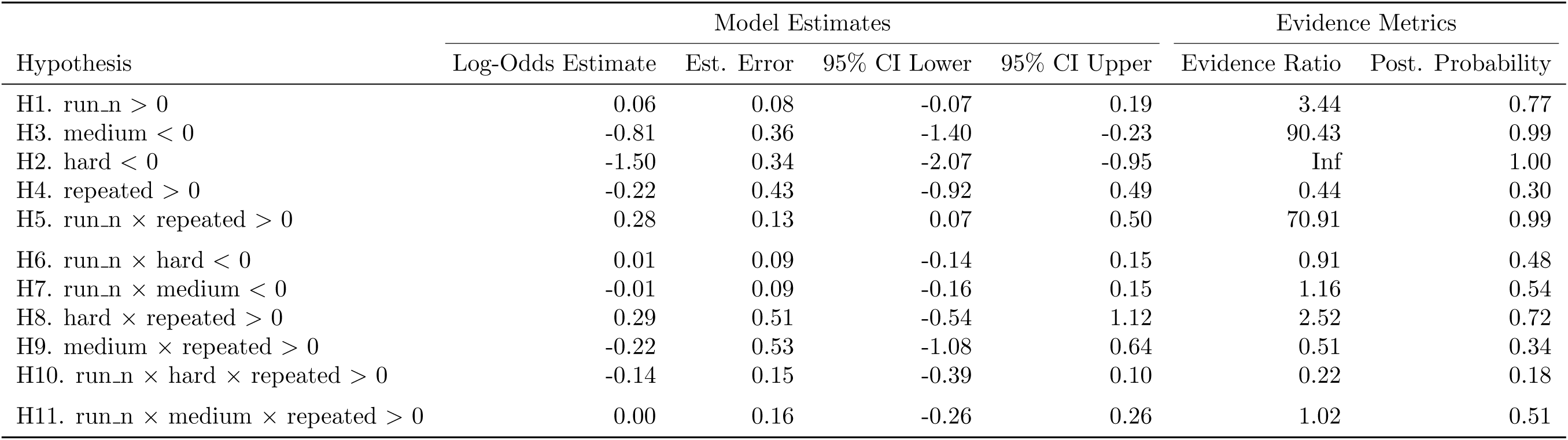
Bayesian Hypothesis Test Results with Practical Significance Estimates.

### B.2 Decoding Results

**Figure S5:**
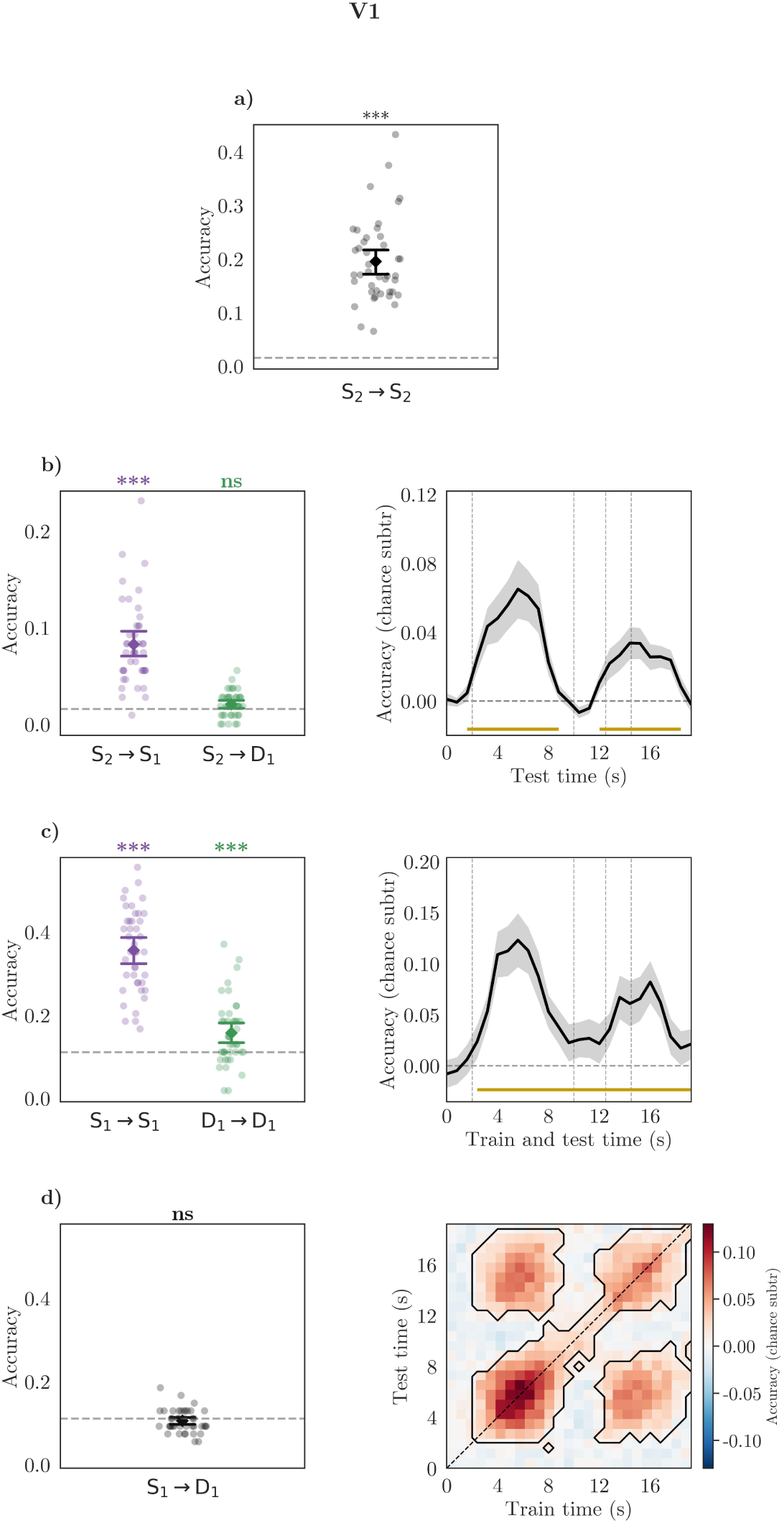
Decoding results for ROI *V1*.

**Figure S6:**
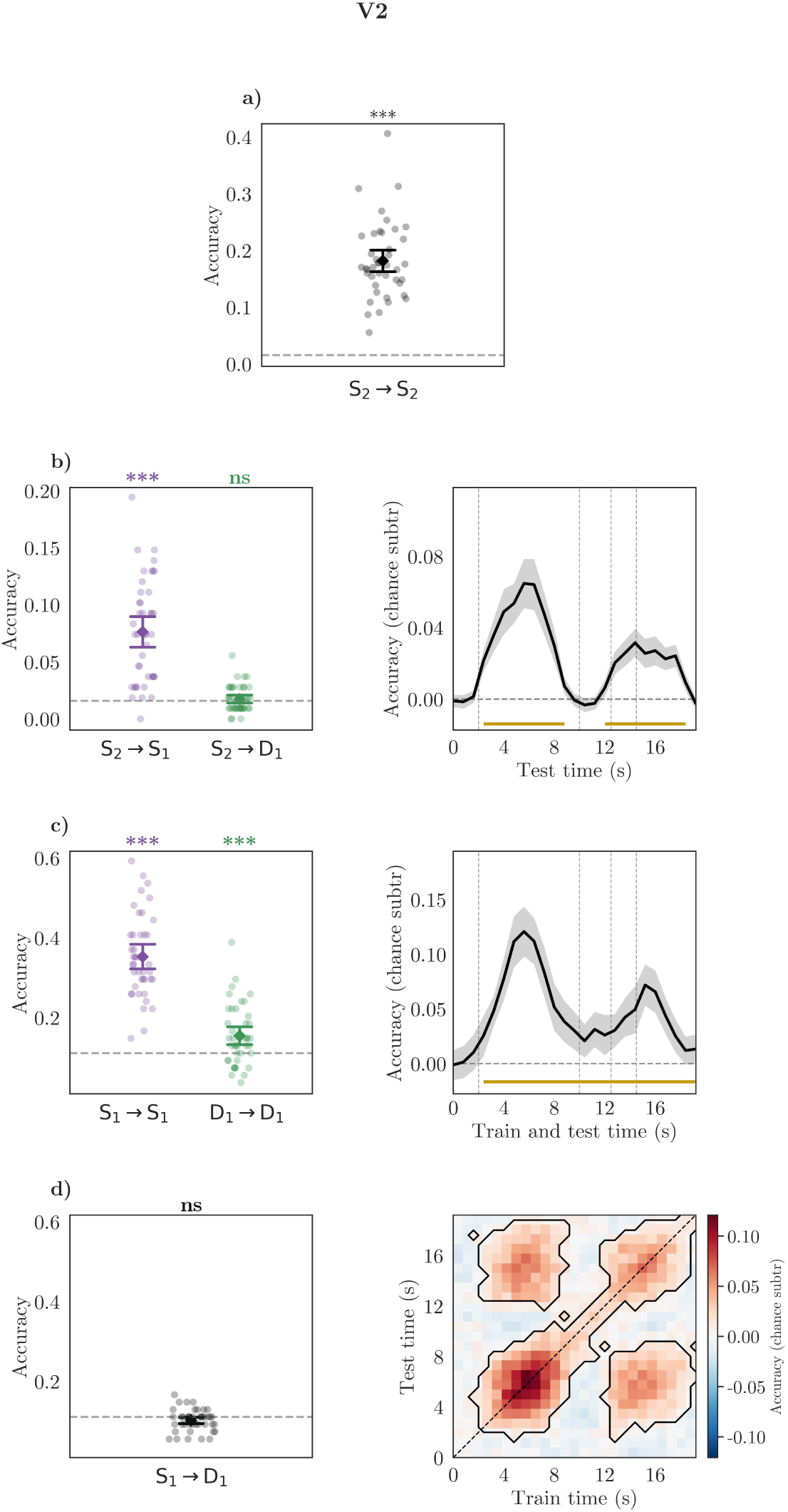
Decoding results for ROI *V2*.

**Figure S7:**
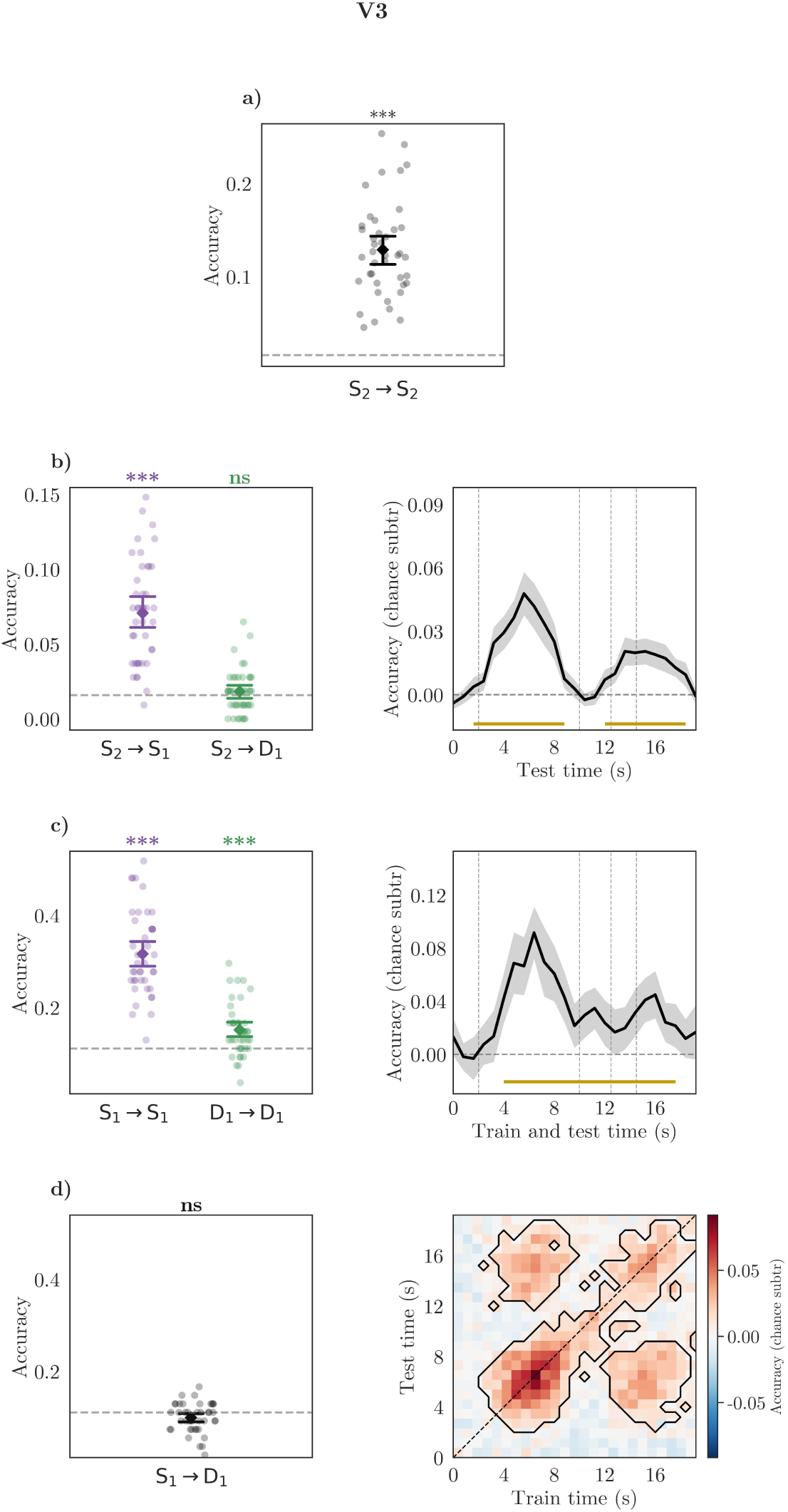
Decoding results for ROI *V3*.

**Figure S8:**
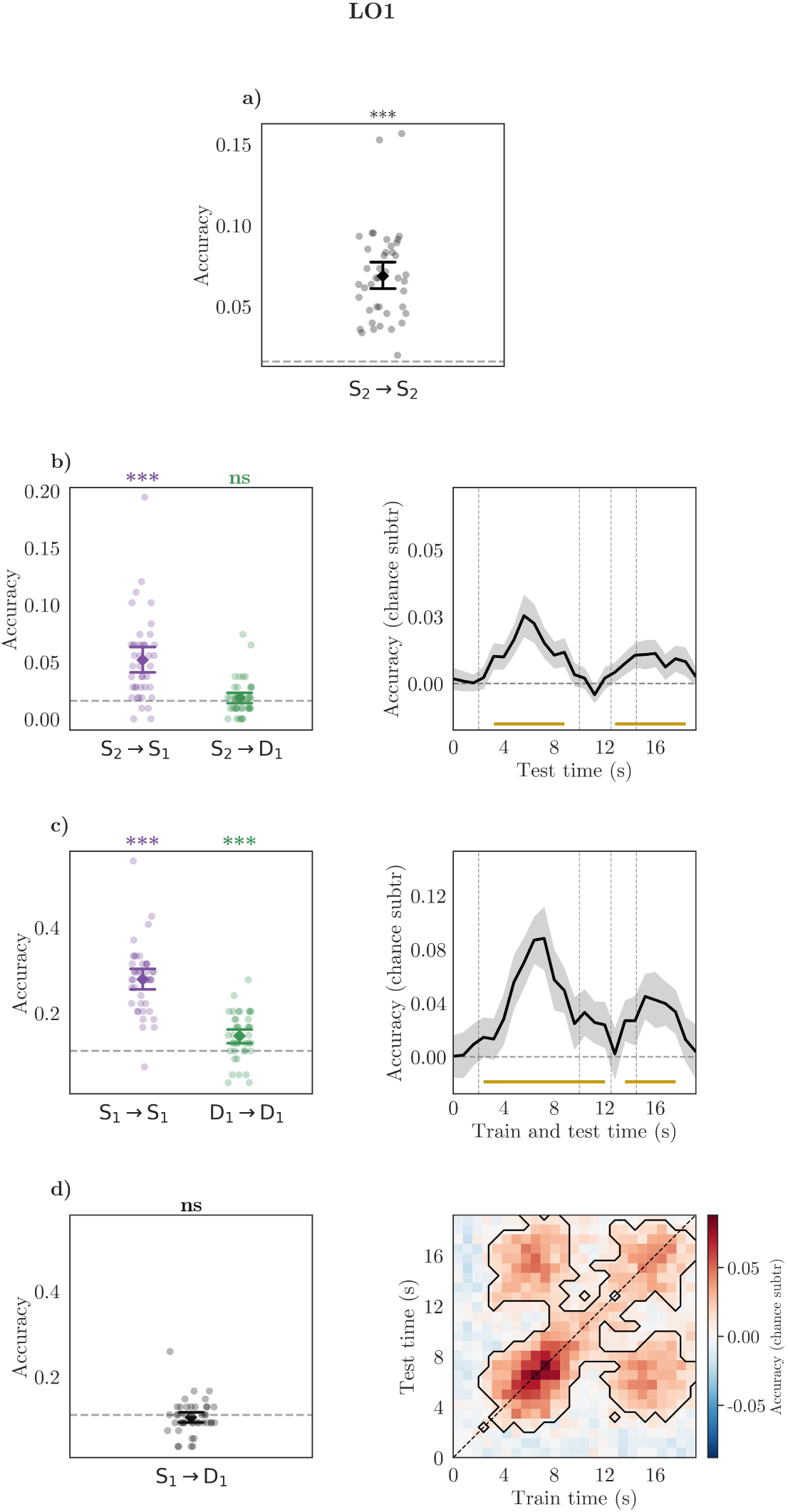
Decoding results for ROI *LO1*.

**Figure S9:**
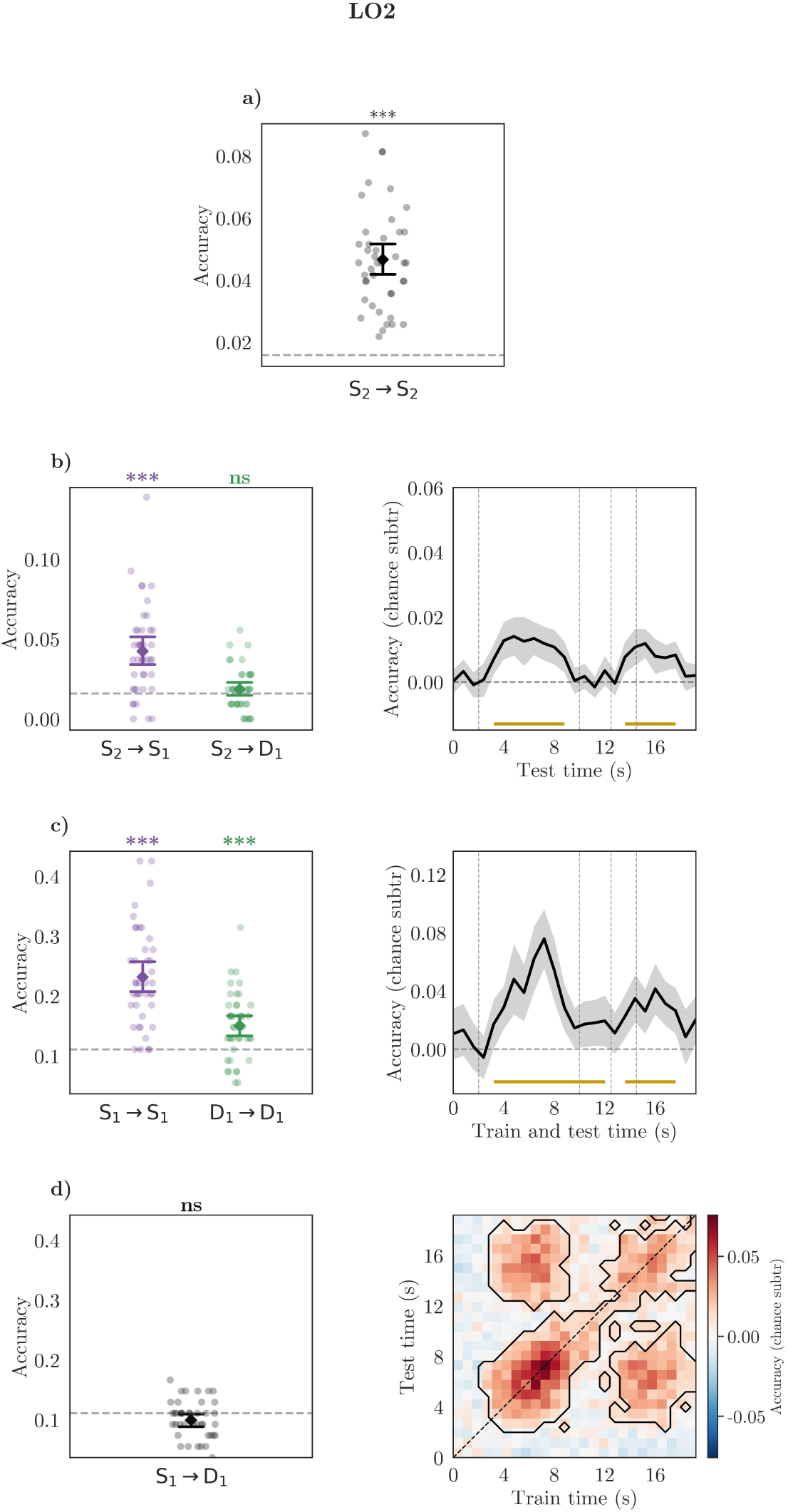
Decoding results for ROI *LO2*.

**Figure S10:**
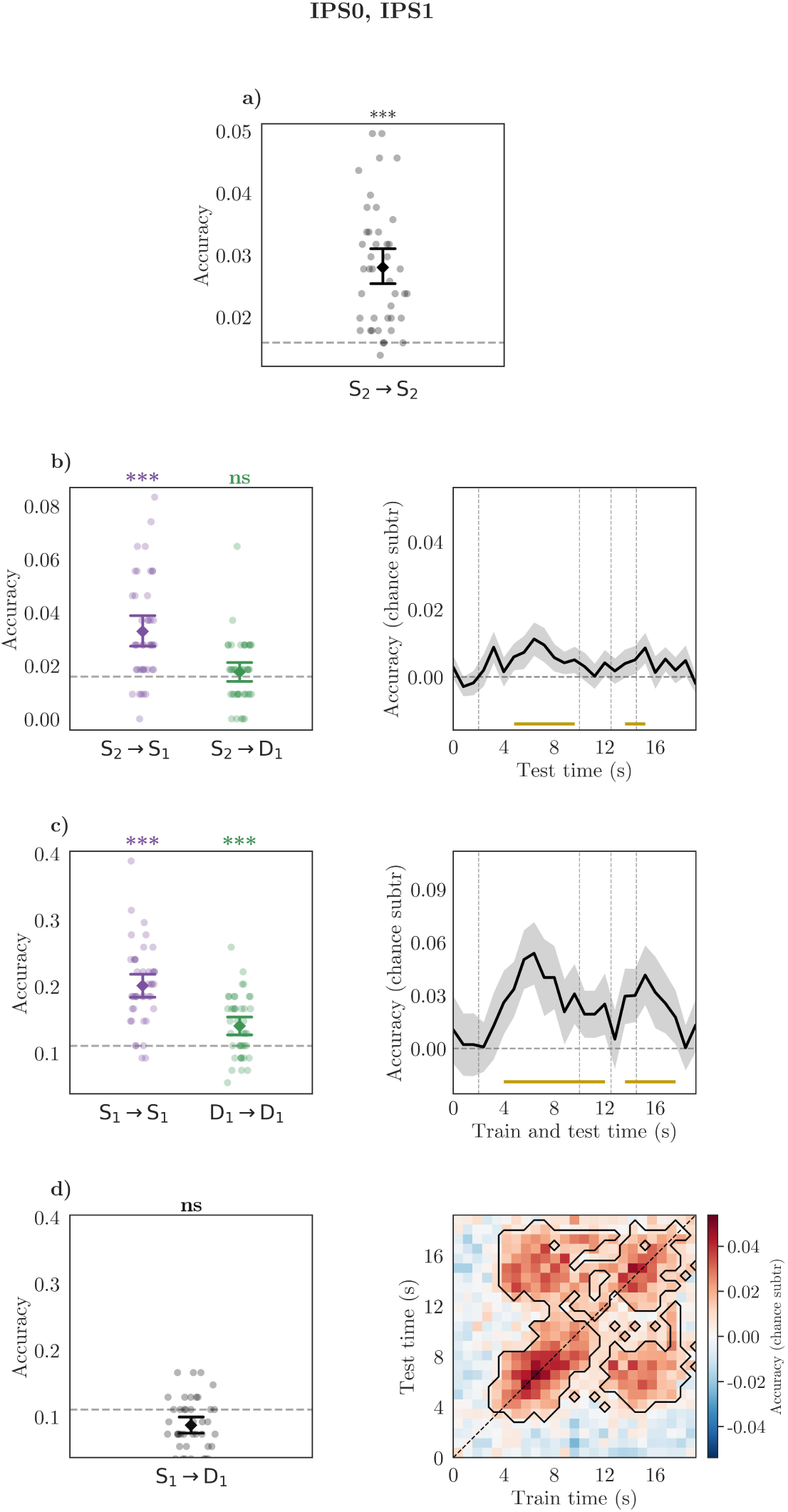
Decoding results for ROI *IPS0IPS1*.

**Figure S11:**
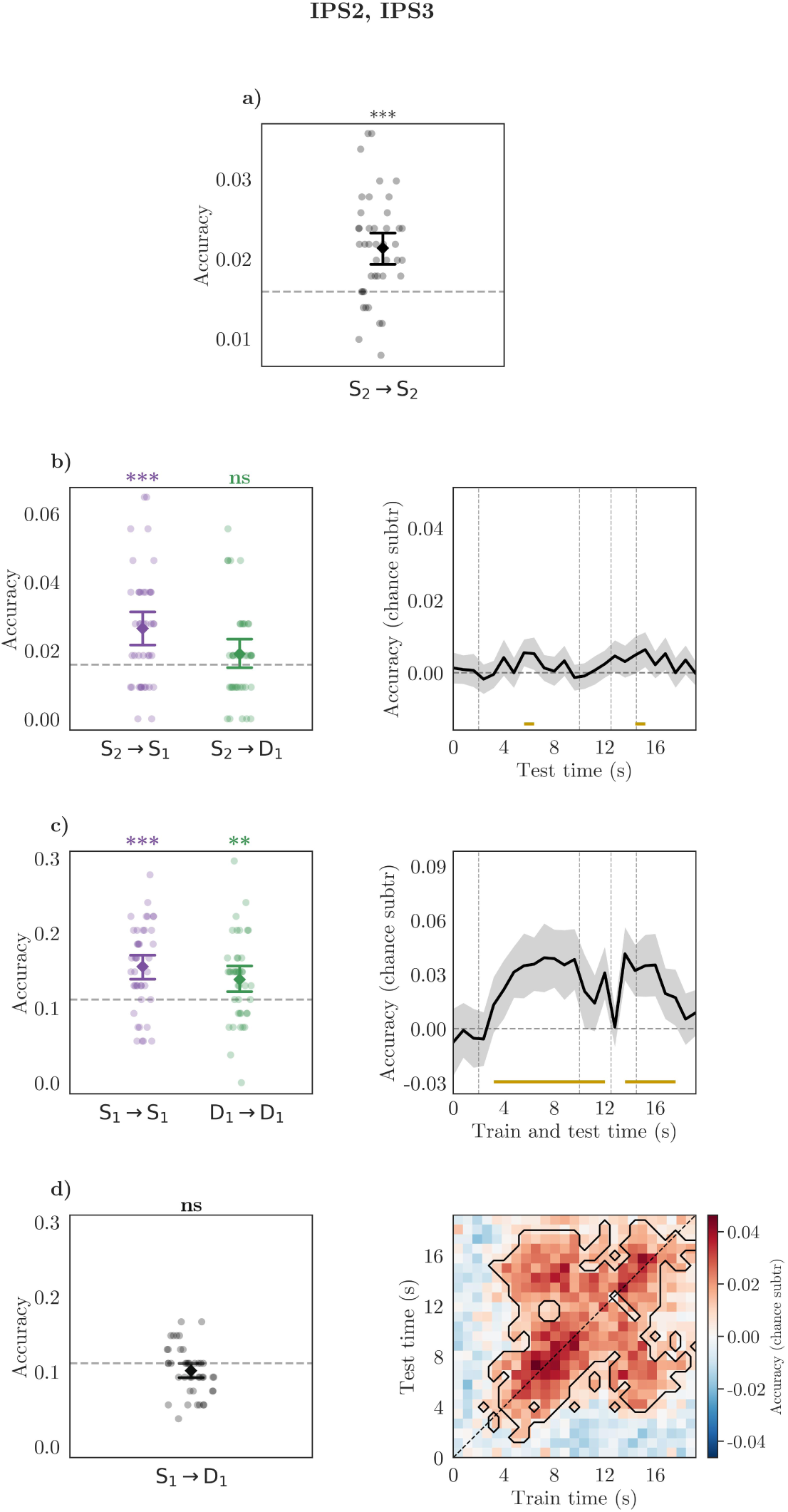
Decoding results for ROI *IPS2IPS3*.

**Figure S12:**
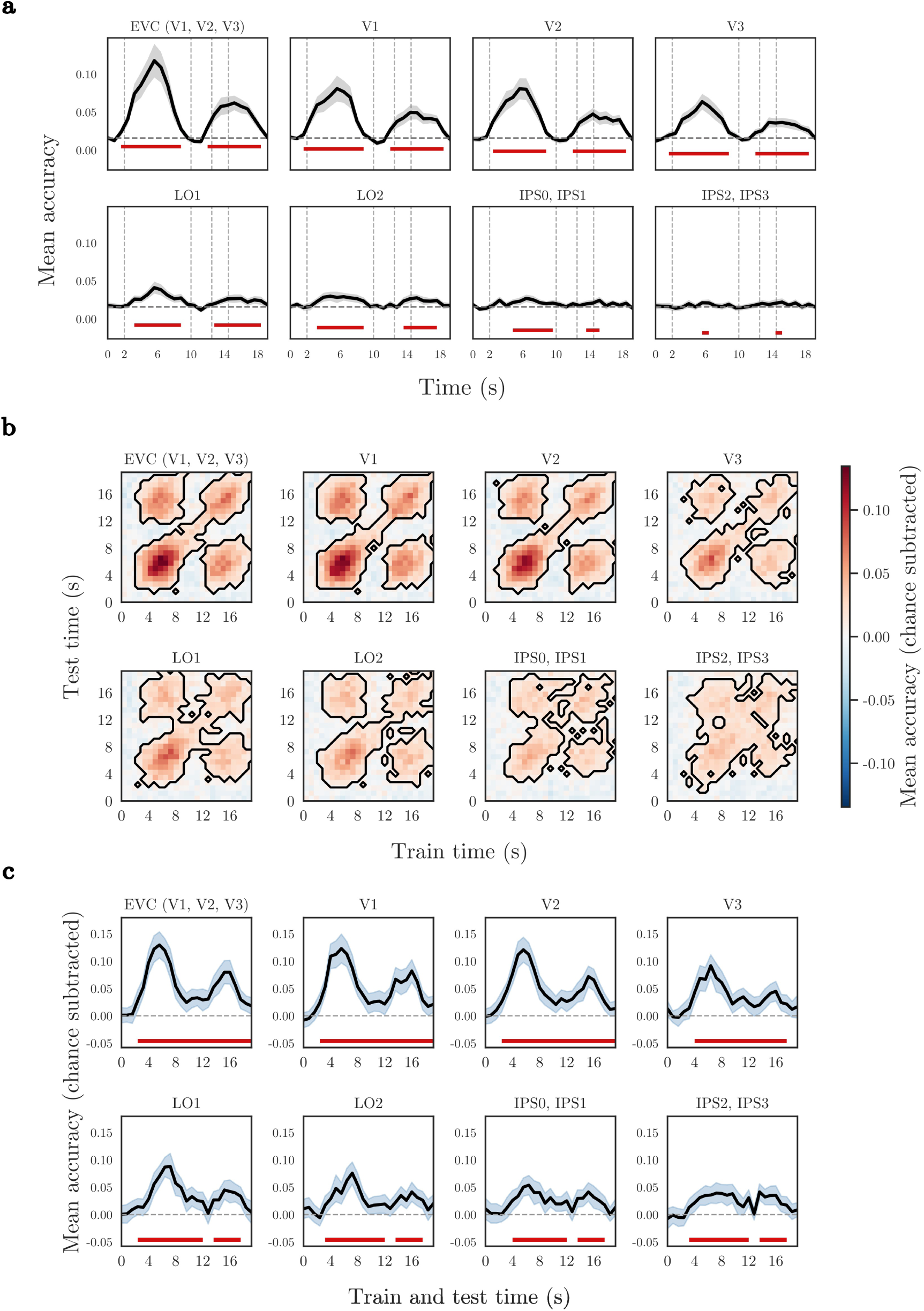
Comparative view of time-resolved classification and temporal generalisation performance across visual and parietal ROIs using raw voxel time series. (a) Cross-session classification accuracy over time. Classifiers were trained on data from Session 2, using a fixed time window from 4 to 8 seconds after stimulus onset (see Methods), and tested on data from Session 1 across the first 25 TRs (20 s) following stimulation onset (time 0 s). Thick black curves show the participant mean (N = 42), the grey bands are 95% CIs. The dashed horizontal line marks chance (63 classes, ≈1.6%). Red bars along the x-axis highlight epochs where classification accuracy was significantly above chance, as determined by a one-sided, sign-flip cluster-mass permutation test (10,000 permutations, cluster-defining threshold *p <* 0.05). Vertical dashed lines mark task phases: delay onset (2 s), probe presentation (10 s), response window onset (12.5 s), and end (14.5 s). The plot extends 5 seconds beyond the trial end to capture the haemodynamic response. (b) Within-session temporal generalisation (Session 1). Classifiers were trained and tested on repeated trials from Session 1 across a 25 TR (20 s) time window following stimulus onset. Heat maps show group mean classification accuracies. Significant clusters, outlined with black contours, were determined using a maximum cluster-mass sign-flip permutation procedure. (c) Within-time decoding. Mean classification accuracy over time with 95% confidence intervals, obtained when classifiers were trained and tested at the same timepoint. Chance level was subtracted before averaging across 42 participants. The dashed horizontal line indicates chance-level classification. Significant values from the cluster-mass permutation test are underscored with red bars (see Methods, Section 3.6.6).

### B.3 Correlation-based Decoder

**Figure S13:**
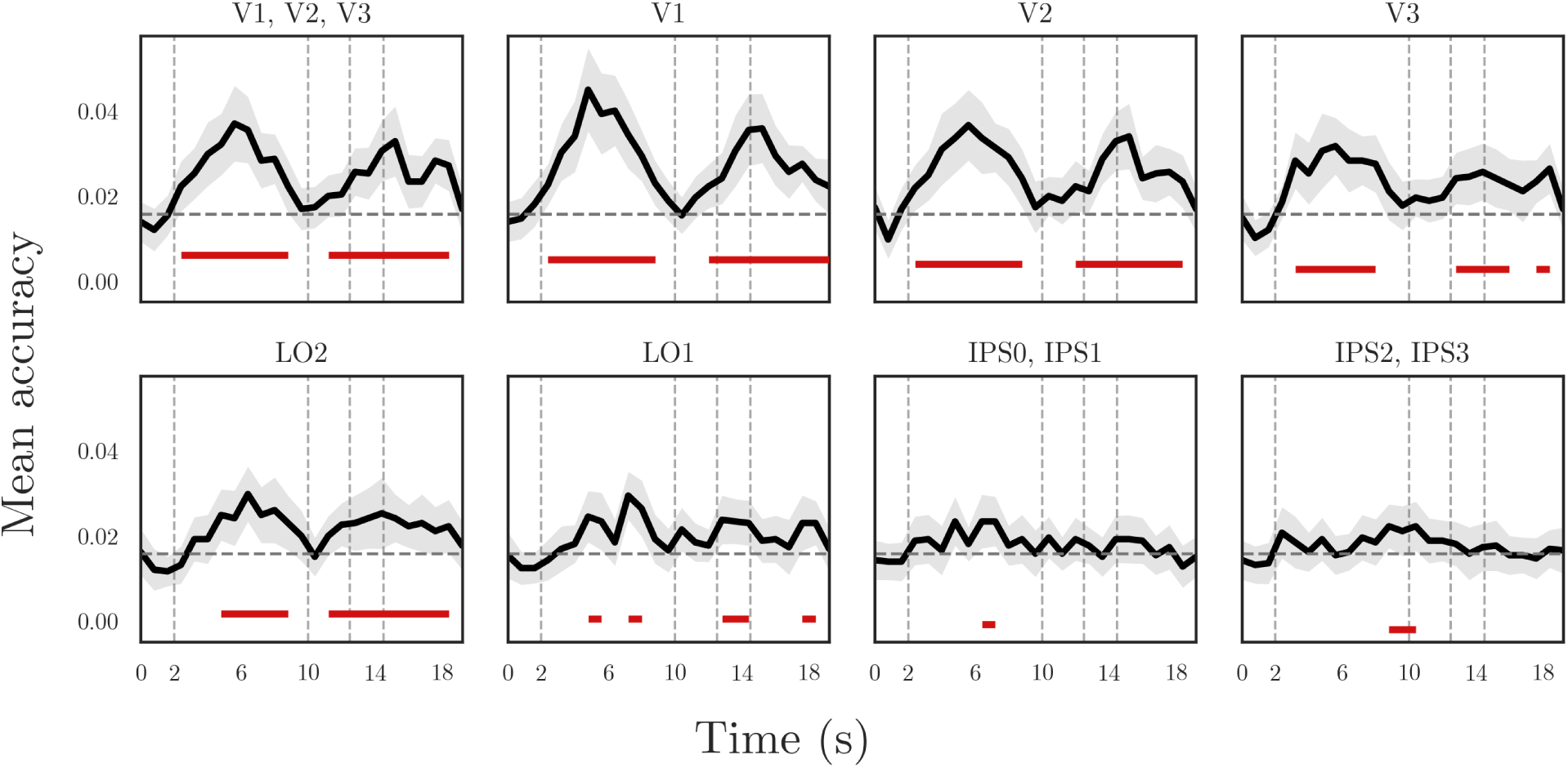
Time-resolved decoding using the correlation-based classifier, which used object-scene-specific patterns from Session 2 to decode object-scene identity at each timepoint in Session 1. For each object-scene, we derived one mean activation pattern per label by averaging its 8 activation patterns from Session 2. At each timepoint of Session 1 trials, we computed the Pearson’s correlation coefficient between the multivoxel pattern at that timepoint and all the 63 Session-2 mean activation patters. For each trial and timepoint, we thus obtained one correlation value per label (63 in total). To calculate classifier accuracy, we extracted the label with the highest correlation value at each TR, compared it to the true label of the current trial, and classified matches as correct and mismatches as incorrect. Here, mean accuracy over time is shown for each ROI (black lines), with 95% confidence intervals (grey shading). The dashed horizontal line indicates chance level (63 classes, ≈1.6%). Vertical dashed lines mark the following task events: delay onset (2 s), probe onset (10 s), response window onset (12.5 s), and response offset (14.5 s). Red bars along the x-axis indicate timepoints at which decoding accuracy was significantly above chance, based on a one-sided, sign-flip cluster-mass permutation test (10,000 permutations, cluster-defining threshold *p <* 0.05). Consistently with the classification results, this decoder shows that perception-based patterns from Session 2 fail to generalise throughout the delay phase, while they robustly decode stimulus identity both during target and probe presentation.

### B.4 Searchlight Analyses

**Figure S14:**
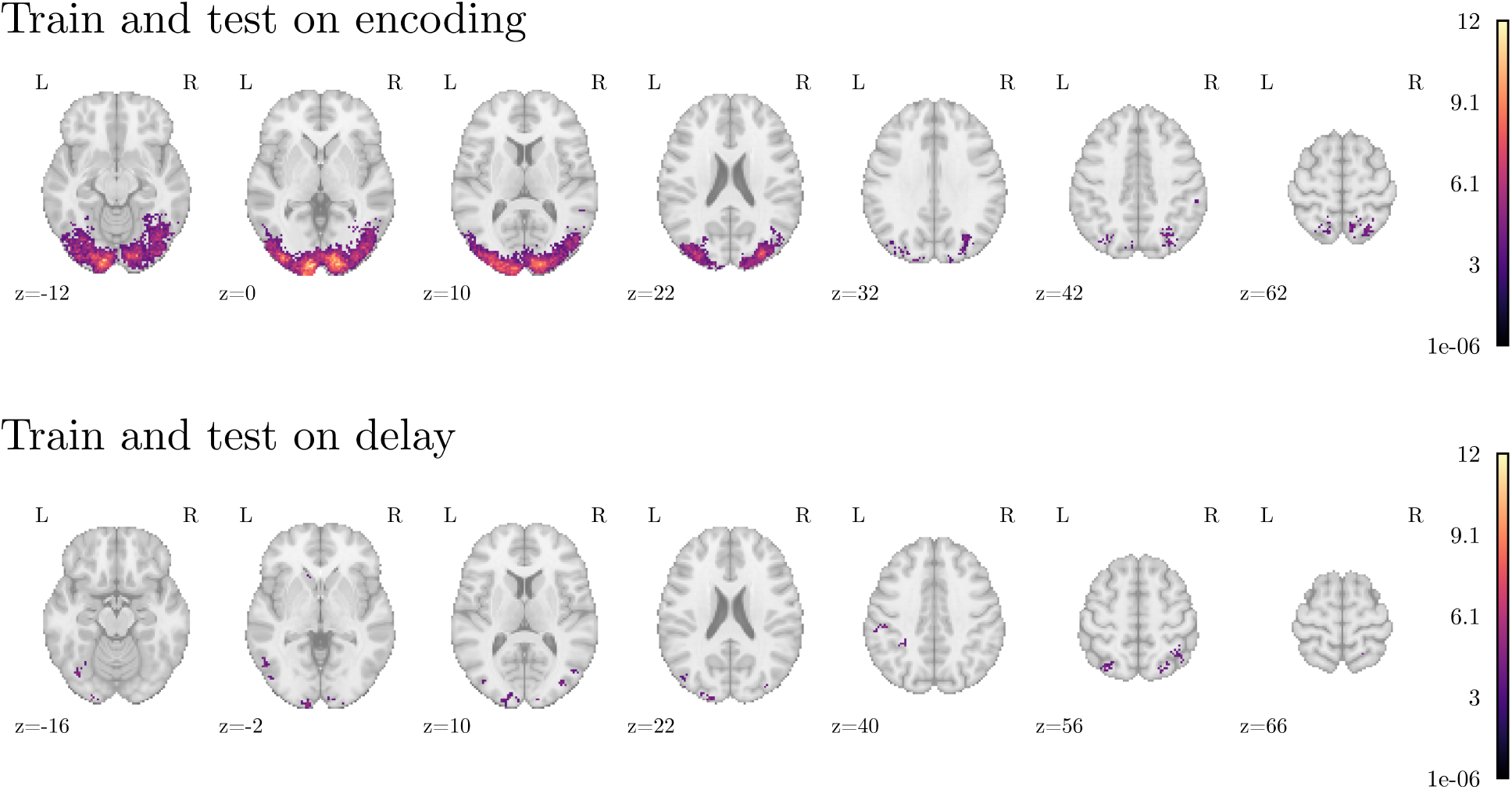
Within-session searchlight results from Session 1. Whole-brain searchlight analysis (sphere radius = 4 mm) was performed using GLM parameter estimates from either the stimulation (top panel) or delay period (bottom panel). Classifier training and testing were conducted within each period. Statistical maps show thresholded z-scores (cluster-corrected; voxelwise *FPR <* 0.001; cluster-extent*>* 13 voxels), overlaid on axial slices in MNI space. Stimulus-specific information could be reliably decoded in visual areas both in the stimuluation and delay period. In the delay period, we observed clusters in the early visual cortex, higher-order object-selective regions, superior parietal lobule and intraparietal sulcus. No significant clusters were found when training on stimulation and testing on delay, or vice versa (results not shown).

**Figure S15:**
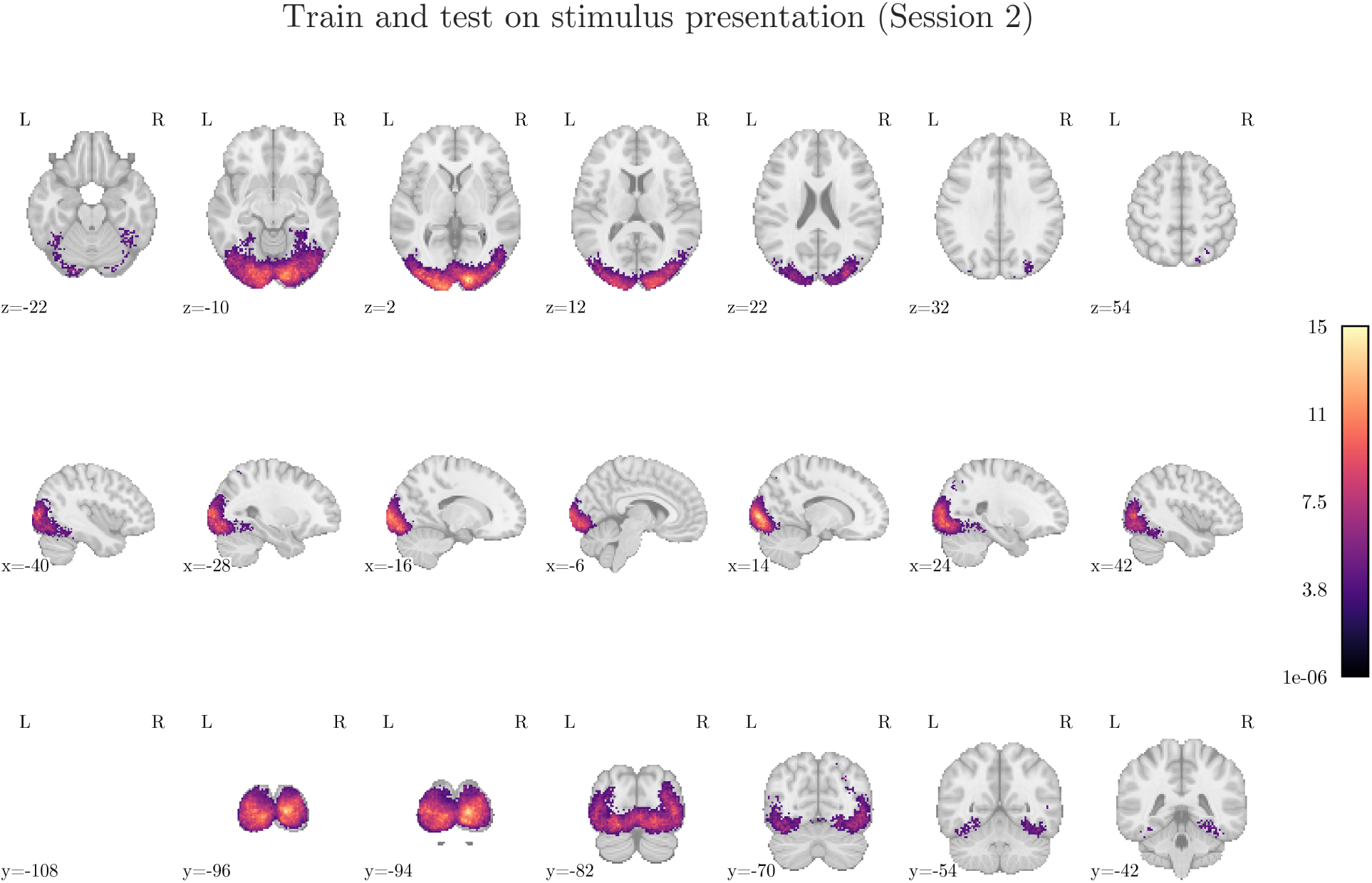
Whole-brain searchlight analysis (sphere radius = 4 mm) was performed training and testing classifiers on GLM parameter estimates from stimulus presentation within Session 2. Thresholded z-maps (cluster-corrected, *FPR <* 0.001; *clusters >* 9*voxels*) are displayed on axial, sagittal, and coronal slices in MNI space. Strong classification was observed in bilateral occipital cortex, consistent with visual stimulus processing.

